# Human iPSC-derived liver organoids model multicellular tissue responses and therapeutic rescue in Wolman disease

**DOI:** 10.64898/2025.12.16.694623

**Authors:** Davide Selvestrel, Caterina Da Rodda, Beatrice Anfuso, Marine Laurent, Annamaria Antona, Alessia Mattivi, Suresh Velnati, Karin Hofmann, Andrea Marfoglia, Rebecca Bertolio, Luciano Conti, Deborah Bonazza, Fabrizio Zanconati, Manuela Mastronardi, Nicolò de Manzini, Natalia Rosso, Claudio Tiribelli, Marcello Manfredi, Daniela Capello, Philippe Drabent, Luca Fava, Silvia Palmisano, Giannino Del Sal, Mario Amendola, Giovanni Sorrentino

## Abstract

Wolman disease (WD), the severe infantile form of lysosomal acid lipase deficiency, is a rare metabolic disorder caused by inactivating mutations in the LIPA gene. Although WD is characterized by profound hepatic dysfunction, experimental human systems capable of modelling multicellular liver pathology and supporting therapeutic testing remain limited. Here, we generated an isogenic human model of WD by introducing LIPA loss-of-function mutations into induced pluripotent stem cells and differentiating them into multicellular human liver organoids (HLO). LIPA-deficient HLO preserved hepatic lineage specification while recapitulating key biochemical and cellular features of WD, including loss of LIPA activity, lysosomal expansion, lipid accumulation, and activation of inflammatory and fibrogenic programs. Single-cell RNA sequencing resolved cell-type–specific disease states across hepatocyte-, stromal-, and biliary-like populations, revealing the emergence of a reactive biliary program consistent with ductular reaction, a complex tissue response associated with chronic liver injury. Importantly, this reactive biliary phenotype was supported by targeted gene-expression analysis in WD liver organoids and independently validated in liver tissue from mouse models and WD patients. Isolated LIPA-deficient cholangiocyte organoids failed to reproduce the DR-associated program, indicating that this response depends on multicellular interactions within the hepatic microenvironment rather than on biliary cell-autonomous dysfunction alone. Consistently, hepatocyte-directed AAV-mediated restoration of LIPA expression attenuated metabolic stress, inflammatory and fibrogenic programs, and suppressed ductular reaction both in organoids and in vivo. Together, these findings establish multicellular human liver organoids as a physiologically relevant platform for modelling emergent tissue-level responses in WD and for evaluating therapeutic rescue strategies in a human context.

## Introduction

Lysosomal acid lipase deficiency (LAL-D) is a rare autosomal recessive lysosomal storage disorder caused by biallelic mutations in the *LIPA* gene, encoding lysosomal acid lipase (LAL), the only enzyme capable of hydrolysing cholesteryl esters and triglycerides within the lysosome^1^. Loss of LAL activity leads to massive accumulation of neutral lipids in hepatocytes, macrophages, and other cell types, resulting in hepatosplenomegaly, dyslipidaemia, liver fibrosis, and multiorgan failure^2^. The clinical presentation of LAL-D spans a spectrum: the most severe form, Wolman disease (WD), manifests within the first weeks of life and is typically fatal in infancy due to liver failure, adrenal calcification, and malabsorption^3^. In contrast, cholesteryl ester storage disease (CESD) presents later in childhood or adolescence with partial residual LAL activity and a slower progression toward hepatic fibrosis, often misdiagnosed as metabolic associated steatotic liver disease (MASLD) or familial hypercholesterolemia^3^.

Current treatments for WD are unfortunately limited. Enzyme replacement therapy (ERT) with Sebelipase alfa has shown clinical benefit and is approved for both WD and CESD^4–6^. However, ERT does not fully prevent disease progression, requires lifelong administration, and may be limited by immune responses and poor penetration in certain tissues^4–6^. More recently, gene therapy approaches using adeno-associated viral (AAV) vectors have shown promise in preclinical models of WD, but no such approaches have been applied in human *in vitro* platform to screen vectors or validate efficacy prior to clinical assessment^7,8^. Moreover, despite extensive characterization in animal models, the molecular and cellular mechanisms driving disease progression in WD livers remain poorly understood, largely due to the absence of multicellular human systems capable of capturing the complex interactions among parenchymal and stromal compartments. These gaps underscore the urgent need for human models that faithfully recapitulate *in vitro* the hepatic cellular, metabolic, and molecular features of WD. In this context, human induced pluripotent stem cell (hiPSC)-derived liver organoids have recently emerged as a powerful tool to model monogenic liver diseases, providing a 3D microenvironment with functional hepatocytes, cholangiocytes, and stromal lineages^9–12^. These systems have been successfully used to study disorders such as alpha-1 antitrypsin deficiency, citrullinemia, and Alagille syndrome, demonstrating their capacity to model complex disease-relevant phenotypes and test gene or drug-based therapies^13–15^. Although *in vitro* models of WD, derived from reprogramming of patient-derived fibroblasts, have been reported, they rely on scarce and hard-to-access primary cells, limiting their reproducibility and widespread use^10^. Moreover, such models inherently carry patient-specific genetic backgrounds, complicating the interpretation of disease phenotypes. These limitations underscore the need for standardized, genetically engineered models derived from widely available human cell lines, which would provide broad accessibility, enable isogenic comparisons, and overcome variability inherent to patient-derived material^16^. Beyond that, WD affects multiple hepatic cell types in distinct yet interconnected ways, including parenchymal and non-parenchymal cells^17,18^. Consequently, models relying on homogeneous cell populations or bulk analyses may fail to capture key cell-type–specific alterations and the microenvironmental interactions that orchestrate disease progression. Therefore, multicellular human liver models are essential to resolve the complex cellular cross-talk and lineage-specific pathophysiology underlying WD^19^.

In this study, we established a human model of WD by CRISPR/Cas9 engineering of a commercial hiPSC line to generate isogenic *LIPA*-deficient pairs and differentiating them into multicellular HLO. Using biochemical assays, lipidomics, and single-cell RNA sequencing, we defined the cell type–specific consequences of *LIPA* loss across hepatocyte, cholangiocyte, and stromal compartments. Notably, this analysis revealed the emergence of a reactive biliary program consistent with ductular reaction (DR), a complex tissue response associated with chronic liver injury, which was further mirrored in livers from WD mice and patients. Importantly, *LIPA*-deficient biliary organoids failed to reproduce this phenotype, supporting the notion that DR in WD depends on multicellular interactions within the hepatic microenvironment rather than on biliary cell-autonomous dysfunctions. Consistently, hepatocyte-directed restoration of LIPA expression suppressed DR both in vivo and in organoids, indicating that correction of parenchymal dysfunction is sufficient to attenuate higher-order reactive tissue responses. Finally, by restoring LIPA specifically in hepatocytes through AAV-mediated delivery in both *Lipa* knockout (*Lipa^-/-^*) mice and WD organoids, we show that therapeutic correction ameliorates disease-associated molecular and cellular phenotypes and suppresses DR, without evidence that LIPA deficiency primarily impairs hepatic lineage differentiation. Together, these findings establish multicellular human liver organoids as a scalable and physiologically relevant platform to model emergent tissue-level responses in WD and to evaluate therapeutic rescue strategies in a human context.

## Results

### Characterization of genetically engineered *LIPA*-deficient human iPSCs

To establish a human *in vitro* model of WD, we generated isogenic *LIPA^-/-^* human induced pluripotent stem cells (hiPSCs) using a commercially available line and CRISPR/Cas9 technology^20^. The targeting strategy involved a single guide RNA designed to introduce a frameshift-inducing nonsense mutation within the *LIPA* coding sequence (Figure 1A). Genotyping of the edited cells revealed efficient mutagenesis at the targeted locus, with a high proportion of frameshift-inducing indels (Figure 1B, Supplementary Figure 1A). Immunofluorescence confirmed the complete loss of LAL protein in lysosomes of *LIPA^-/-^* cells (Figure 1C). Western blot analysis independently confirmed the complete absence of LAL protein in *LIPA^-/-^* cells (Supplementary Figure 1B), and gene expression analysis showed a dramatic reduction in *LIPA* mRNA levels, consistent with degradation via nonsense-mediated decay (Figure 1D).

**Fig. 1:**
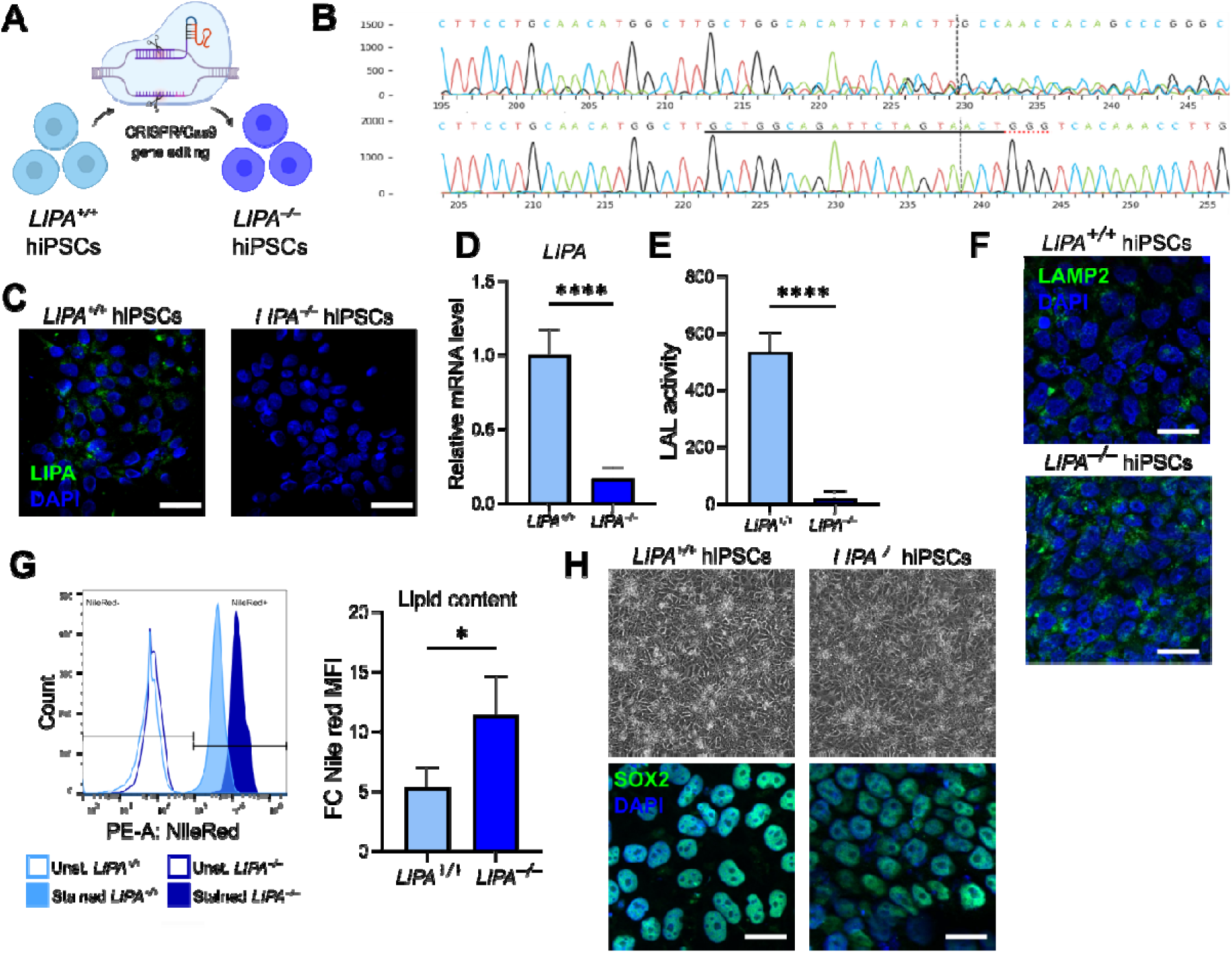
Characterization of genetically engineered *LIPA*-deficient hiPSCs. **A** Schematic representation of CRISPR/Cas9-mediated knockout of the LIPA gene in human induced pluripotent stem cells (hiPSCs). **B** Representative Sanger sequencing chromatograms confirming the successful editing of the *LIPA* locus in hiPSCs. The black line denotes the gRNA binding site, the red dotted line the PAM sequence, and the black dotted line the predicted Cas9 cleavage site. **C** Immunofluorescence (IF) analysis showing complete loss of LIPA protein in *LIPA^-/-^* hiPSCs. **D** Quantitative RT–PCR analysis of LIPA mRNA expression levels in *LIPA^+/+^* and *LIPA^-/-^* hiPSCs. Results represent mean ± SD; n = 3. \*\*\*\**p*□≤□0.0001 Student’s t-test was applied. **E** Lysosomal acid lipase (LAL) enzymatic activity in *LIPA^+/+^* and *LIPA^-/-^* hiPSCs. Results represent mean ± SD; n = 4. \*\*\*\**p*□≤□0.0001 Student’s t-test was applied. **F** Immunofluorescence staining for LAMP2 (green) showing lysosomal distribution in *LIPA^+/+^*and *LIPA^-/-^* hiPSCs. Nuclei were stained with DAPI (blue). Scale bars: 50 μm. **G** Flow cytometric analysis of Nile Red staining (left panel) and quantification (right panel) of Nile Red mean fluorescence intensity (MFI), expressed as fold change relative to the mean value of *LIPA*^+/+^ hiPSCs, showing increased neutral lipid accumulation in *LIPA*^-/-^ hiPSCs. Results represent mean ± SD; n = 3. Student’s t-test was applied: * *p*□≤□0.05. **H** Bright-field images indicate no morphological differences between *LIPA^+/+^* and *LIPA^-/-^* hiPSCs (top panels) and immunostaining for SOX2 (green) confirms preserved pluripotency in both genotypes (bottom panels). Nuclei were counterstained with DAPI (blue). Scale bars: 50 μm.

To confirm functional ablation of lysosomal acid lipase activity, we performed enzymatic assays on *LIPA^+/+^* and *LIPA^-/-^* iPSCs. *LIPA^-/-^* cells exhibited a complete loss of LAL activity, supporting full knockout efficiency (Figure 1E). In line with the expected lysosomal dysfunction in LAL deficiency, immunofluorescence staining for LAMP2– a marker of lysosomal membranes commonly used to assess lysosomal abundance and morphology– revealed marked expansion of the lysosomal compartment in *LIPA^-/-^* cells as compared to controls (Figure 1F)^21^. To further corroborate the characteristic phenotype associated with WD, we quantified intracellular neutral lipid accumulation by Nile Red staining and flow cytometry, revealing a marked increase in lipid content in *LIPA^-/-^* iPSCs compared to controls (Figure 1G)^22–24^. Notably, *LIPA^-/-^* hiPSCs morphology was similar to *LIPA^+/+^* cells (Figure 1H, upper panel), and expression of pluripotency markers (e.g., OCT4, SOX2) was comparable between *LIPA^+/+^* and *LIPA^-/-^* lines indicating no alteration in pluripotent stem cell identity (Figures 1H, lower panel; Supplementary Figure 1C).

Together, these data validate the successful generation of functional *LIPA*-deficient hiPSC lines that exhibit biochemical and cellular hallmarks of LAL deficiency while preserving pluripotency markers, thus establishing a resource for downstream disease modelling.

### *LIPA* deficiency does not affect proper liver organoid derivation and differentiation

Human liver organoids (HLOs) provide a physiologically relevant system to study liver function and pathology, as they incorporate multiple hepatic lineages and preserve aspects of tissue architecture not captured in standard cultures^10–12^. To model the multicellular hepatic environment affected in WD, we differentiated *LIPA^+/+^* and *LIPA^-/-^* hiPSC lines into HLO following an established protocol^10–12^ which involves a sequential induction of definitive endoderm and foregut, followed by hepatic specification and maturation in 3D, embedded in a basement membrane extract (BME) (Figure 2A)^10–12^. After 21 days of culture, both *LIPA^+/+^* and *LIPA^-/-^*hiPSCs reproducibly formed compact organoids, with similar morphology and size between genotypes (Figure 2B, 2C and Supplementary Figure 2A), in which the epithelial compartment was surrounded by stromal cells, as shown by Vimentin (VIM) and E-cadherin (E-CAD) staining respectively (Figure 2D).

**Fig. 2:**
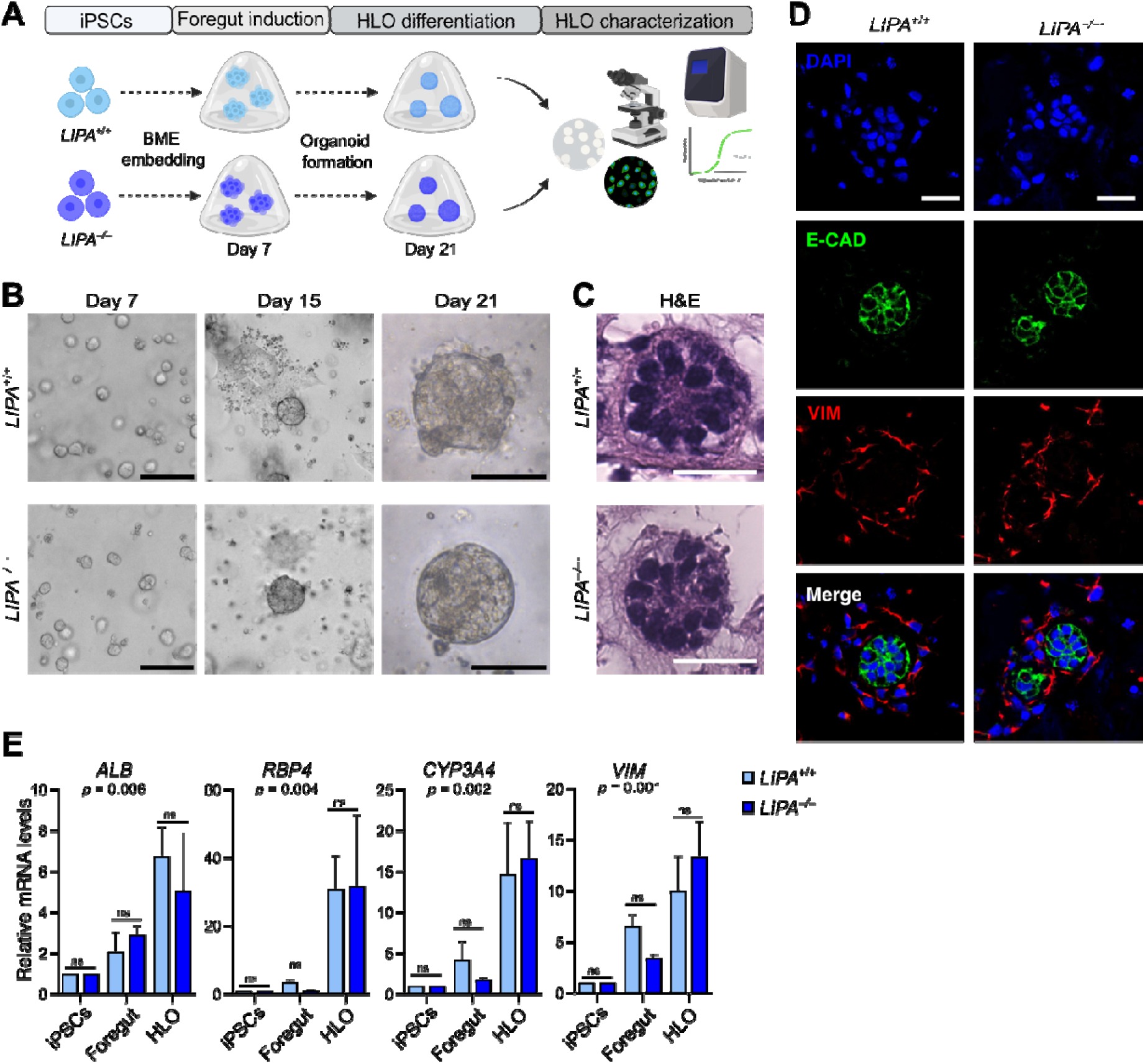
***LIPA* deficiency does not affect proper liver organoid differentiation. A** Schematic representation of the HLO induction protocol. **B** Bright-field image of BME drop containing *LIPA^+/+^* and *LIPA^-/-^* HLO ad day 7, 15 and 21. The scale bar represents 50 μm. **C** Images of haematoxylin and eosin staining in HLO. The scale bar represents 50 μm. **D** Whole-mount immunofluorescent staining of E-cadherin (green), vimentin (red), and nuclei (blue) in HLO at day 21. The scale bar represents 25 μm. **E** Quantitative RT–PCR analysis of differentiation markers in *LIPA^+/+^* and *LIPA^-/-^* cells at day 0 (iPSCs), 7 (foregut) and 21 (HLO). Results represent mean ± SD; n = 4-6 independent differentiation experiments. Two-way ANOVA was applied followed by Tukey post-hoc test. ns = not significant.

In both *LIPA^+/+^* and *LIPA^-/-^* organoids, pluripotency genes such as *OCT4* and *SOX2* were efficiently silenced upon differentiation, confirming proper exit from the stem cell state (Supplementary Figure 2B)^25^. To explore whether *LIPA* deficiency affected organoid differentiation, we measured expression of a panel of marker genes representative of different hepatic populations. Importantly, key hepatocyte markers, such as albumin (*ALB*), the cytochrome *CYP3A4* and retinol binding protein 4 (*RBP4*), were significantly upregulated upon differentiation in both lines^26^ (Figure 2E). Similarly, the stromal marker *VIM* was induced in both *LIPA^+/+^* and *LIPA^-/-^*organoids, suggesting that both parenchymal and mesenchymal differentiation occurred efficiently regardless of *LIPA* status.

Altogether, these data demonstrate that *LIPA*-deficient hiPSCs retain the capacity to form structured, multicellular liver organoids and that *LIPA* deficiency does not impair HLO maturation, allowing hepatic lineages to emerge efficiently.

### *LIPA-*deficiency in HLO drives lipid metabolic rewiring and triggers inflammatory and fibrogenic programs

To assess the molecular impact of *LIPA* deficiency on lipid metabolism in HLO, we first monitored the expression of genes controlling key lipid metabolism pathways in independent differentiation experiments. We observed robust upregulation of Diacylglycerol O-acyltransferase (DGAT1 and DGAT2), which catalyse triglyceride synthesis, as well as 3-Hydroxy-3-Methylglutaryl-CoA Reductase (HMGCR), the rate-limiting enzyme in cholesterol biosynthesis, and Acetyl-CoA Carboxylase Alpha (ACACA), the rate-limiting enzyme in fatty acid biosynthesis (Figure 3A)^27,28^. The induction of these genes is expected and reflects a compensatory cellular response to the lack of accessible lipids caused by defective lysosomal hydrolysis. To corroborate these findings at the metabolite level, we performed lipidomic analysis on *LIPA^+/+^* and *LIPA^-/-^*HLO (Figure 3B). Supervised multivariate analysis revealed a clear separation between two genotypes reflecting profoundly distinct lipidomic signatures, consistent with the strong metabolic rewiring induced by *LIPA* deficiency (Supplementary Figure 3A). Of note, we found a marked upregulation of cholesteryl esters (CEs) and triglycerides in *LIPA^-/-^* organoids (Figure 3C). Moreover, enrichment analysis highlighted significant accumulation of steryl esters, sterols, ceramides, and broader sphingolipid classes, as well as signatures linked to ER and endo-lysosomal membranes (Figure 3D). This pattern is highly consistent with WD pathophysiology and functionally confirms the changes observed in gene expression^24,29,30^.

**Fig. 3:**
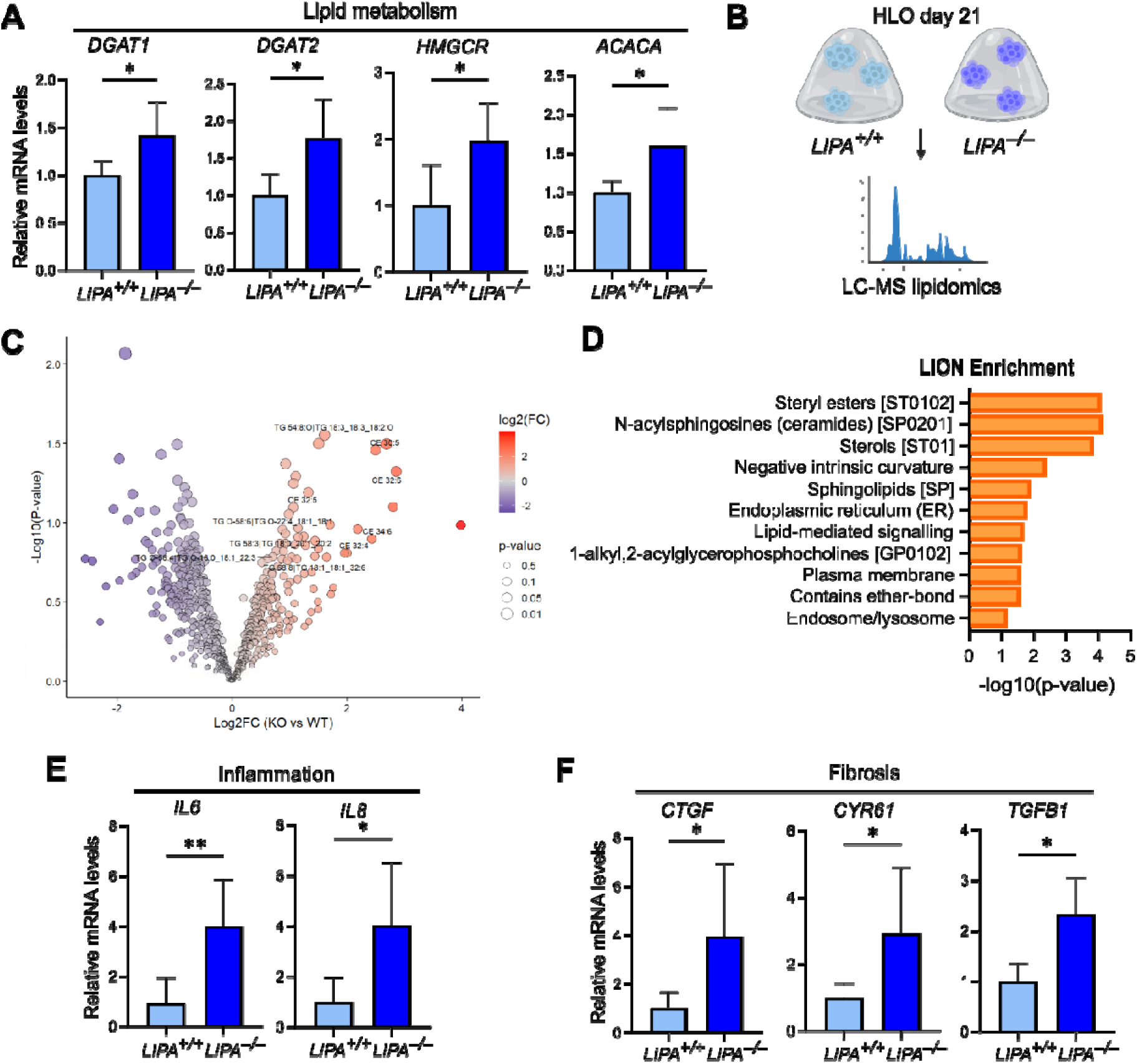
***LIPA-*deficiency in liver organoids drives lipid metabolic rewiring and triggers inflammatory and fibrogenic programs. A** Quantitative RT–PCR analysis of lipid metabolism genes in *LIPA^+/+^* and *LIPA^-/-^* HLOs. Results represent mean ± SD; n = 4-6 independent differentiation experiments. Student’s t-test was applied: * *p*□≤□0.05. **B** Schematic representation of the metabolism-related analyses conducted on *LIPA^+/+^* and *LIPA^-/-^* HLO. **C** Volcano plot showing lipids differentially abundant in *LIPA^-/-^* vs. *LIPA^+/+^* HLO. **D** Lion enrichment analysis on lipids exhibiting differential abundance. **E** Quantitative RT–PCR analysis of inflammation markers in *LIPA^+/+^* and *LIPA^-/-^* HLOs. Results represent mean ± SD; n = 4-6 independent differentiation experiments. Student’s t-test was applied: * *p*□≤□0.05, ** *p*□≤□0.01. **F** Quantitative RT–PCR analysis of fibrosis markers in *LIPA^+/+^* and *LIPA^-/-^*HLOs. Results represent mean ± SD; n = 4-6 independent differentiation experiments. Student’s t-test was applied: * *p*□≤□0.05.

Since metabolic stress in WD does not remain confined to lipid accumulation but drives inflammatory signalling and fibrotic responses, we monitored the expression of key inflammatory and fibrosis-associated genes. Interestingly, inflammatory markers such as interleukin (IL) 6 and IL8 were up-regulated in *LIPA^-/-^* organoids^31–33^ (Figure 3E). Accordingly, fibrogenic mediators, including connective tissue growth factor *(CTGF),* cysteine-rich angiogenic inducer 61 (*CYR61)* and transforming growth factor-beta 1 (*TGF*β*1),* were also significantly induced, suggesting activation of profibrotic pathways, a hallmark of WD livers^34–36^ (Figure 3F).

Together, these results demonstrate that *LIPA*-deficient liver organoids (hereinafter referred to as *WD organoids*) undergo a profound reprogramming of their lipid metabolic landscape as well as inflammatory activation, and induction of fibrogenic programs, faithfully modelling key features of WD in a human *in vitro* system.

### Single-cell transcriptomics reveals cell specific effects of *LIPA* deficiency in WD organoids

To dissect the cellular and molecular consequences of *LIPA* deficiency at high resolution, we performed single-cell RNA sequencing (scRNAseq) on differentiated wild type and WD organoids (Figure 4A). As first, pseudo-bulk Gene Set Enrichment Analysis (GSEA) of differentially expressed genes (DEGs) in all cells confirmed that *LIPA*-deficient cells upregulated pathways associated with regeneration (i.e. cell cycle genes), inflammation (i.e. TNFα), and fibrosis (i.e. TGFβ) (Figure 4B). To further validate whether the transcriptional rewiring observed in human organoids mirrors the phenotype observed *in vivo*, we compared DEGs identified in WD organoids with those from livers of *Lipa^-/-^* mice, which recapitulate major pathological features of WD, given the lack of available transcriptomic data from human patients^7^. Strikingly, a substantial fraction of DEGs identified in human organoids overlapped with those dysregulated in the mouse model (Figure 4C) and GSEA on these overlapping genes revealed strong enrichment for processes related to extracellular matrix organization, collagen formation and degradation, lipid and lipoprotein metabolic pathways—all hallmark processes associated with the WD (Figure 4C). This overlap demonstrates that *LIPA* deficiency in HLO elicits a conserved gene expression program that mirrors the complex hepatic alterations observed *in vivo*.

**Fig. 4:**
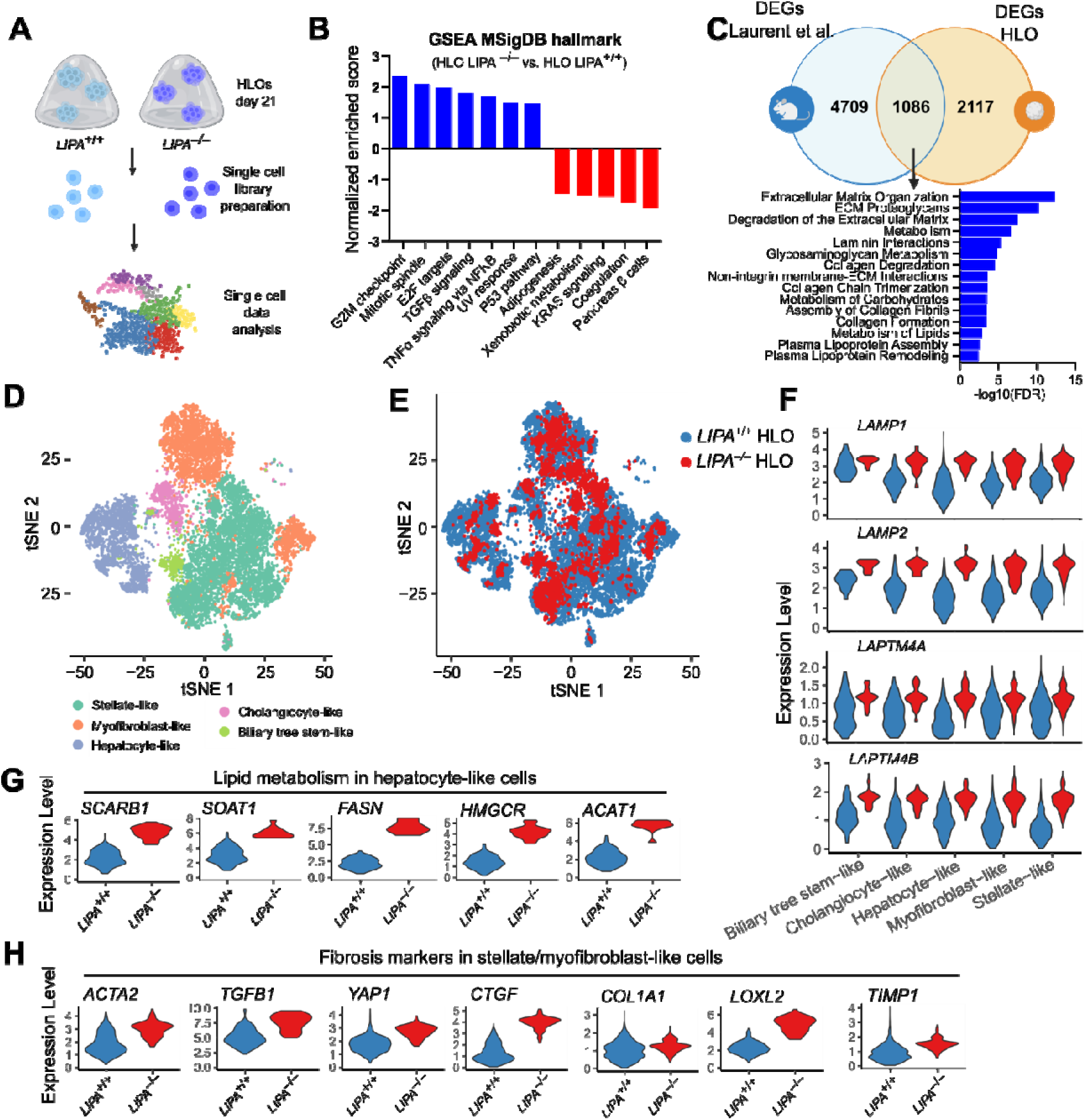
Single-cell transcriptomics reveals cell specific effects of *LIPA* deficiency in WD organoids. **A** Schematic representation of the scRNA-seq experimental workflow. **B** Gene set enrichment analysis (GSEA) conducted in a pseudo-bulk analysis on *LIPA^+/+^* and *LIPA^-/-^* HLO. **C** Venn diagram showing overlap between DEGs from Laurent et al., and the present work (top) and the GSEA conducted on the common genes between the 2 datasets. **D** Overall UMAP showing cell annotation in *LIPA^+/+^* and *LIPA^-/-^* HLOs after quality control. **E** Overall UMAP shown by HLO genotype. **F** Violin plots of lysosome-associated genes in all cell populations grouped by genotype. **G** Violin plots of lipid metabolism genes in hepatocyte-like cells grouped by genotype. **H** Violin plots of fibrosis markers in stellate-like and myofibroblast-like cells grouped by genotype.

We next performed dimensionality reduction and clustering analysis, which revealed the presence of multiple transcriptionally distinct populations in organoid across both genotypes (Figure 4E). Albeit with varying proportions, the overall cellular composition and diversity closely mirrored those previously described by Ouchi et al. for this multicellular liver organoid system, thus confirming the robustness and reproducibility of our organoid model (Supplementary Figure 4A)^10^. Using established lineage markers, we annotated these clusters as hepatocyte-like cells (*TTR, APOA2*), cholangiocyte-like cells (*SPP1, HNF1B*), biliary tree stem-like cells (*NEUROG3, FOXA2*), stellate-like cells (*VIM, DCN*), and myofibroblast-like cells (*TGFBI, CXCL12*) (Figures 4D,E and Supplementary Figure 4B,C)^10^.

To investigate the appearance of a WD phenotype across the different populations, we analysed the expression of lysosomal genes. Consistent with the ubiquitous role of LIPA in lysosomal function, WD organoids showed a broad upregulation of lysosomal membrane proteins, including *LAMP1, LAMP2, LAPTM4A,* and *LAPTM4B* in all cell clusters (Figure 4F). Focusing on hepatocyte-like cells, direct comparison of *LIPA^+/+^*and *LIPA^-/-^* cells revealed the upregulation of key genes involved in cholesterol biosynthesis, trafficking, and storage (Figure 4G). Genes such as ACAT1, HMGCR, SOAT1—components of the cholesterol synthesis and esterification axis—were induced in WD hepatocyte-like cells in response to the perceived shortage of free cholesterol at the ER, a consequence of its sequestration in undegraded lysosomal pools. The lipogenic enzyme fatty acid synthase (FASN), which catalyses de novo fatty acid synthesis, was also upregulated, reflecting a compensatory activation of de novo lipogenesis aimed at replenishing cytosolic lipid pools. In addition, the Scavenger Receptor Class B Member 1 (SCARB1), which mediate cholesteryl esters uptake, weas also upregulated, suggesting an attempt to acquire cholesterol from external sources. Altogether, these changes delineate a maladaptive metabolic program driven by disrupted lysosomal lipid clearance and are in line with the cellular phenotype expected in WD hepatocytes.

Consistent with the multicellular organization of the system, non-parenchymal cells also acquired disease-relevant states in response to *LIPA* loss. Interestingly, in WD organoids we observed a strong activation of the stromal/mesenchymal compartment, characterized by increased expression of fibrogenic markers, including *ACTA2* (α-SMA) and *TGF*β, the mechanoresponsive transcriptional co-activator *YAP1*, and extracellular-matrix genes *COL1A1* and *LOXL2*, together with matricellular regulators such as *CTGF* and *TIMP1* (Figure 4H)^34,37,38^.

Collectively, these results provide the first high-resolution view of WD across multiple hepatic lineages in a human system, highlighting distinct cell-type–specific alterations associated with *LIPA* deficiency.

### HLO capture ductular reactive states emerging in WD

We noticed that the relative abundance of biliary stem-like cells in WD organoids increased, rising from 1.5% to 6.3% (Supplementary Figure 5A). Consistent with this observation, independent experiments of HLO differentiation demonstrated a substantial increase in expression of *KRT19*, a biliary lineage marker^39^ (Figure 5A). This phenotype was consistent with a ductular reaction (DR), a conserved tissue response to chronic liver injury characterized by the expansion and remodeling of biliary epithelial and progenitor-like cells accompanied by hepatocyte-to-biliary lineage reprogramming^40^. Consistently, *LIPA^-/-^* biliary cells displayed a clear upregulation of TNF Receptor Superfamily Member 12A (*TNFRSF12A)* and Neural Cell Adhesion Molecule 1 (*NCAM1)*— two well established markers of DR—together with induction of proliferation genes such as *MKI67* and *TOP2A*, which are hallmarks of the reactive biliary phenotype (Figure 5B)^41,42^ Moreover, YAP1—a key regulator of biliary epithelial plasticity and implicated in hepatocyte-to-biliary reprogramming during liver injury, as well as its transcriptional targets (*CTGF, BIRC5, COL12A1*)– were induced in *LIPA^-/-^* biliary cells (Figure 5B)^43,44^.

**Fig. 5:**
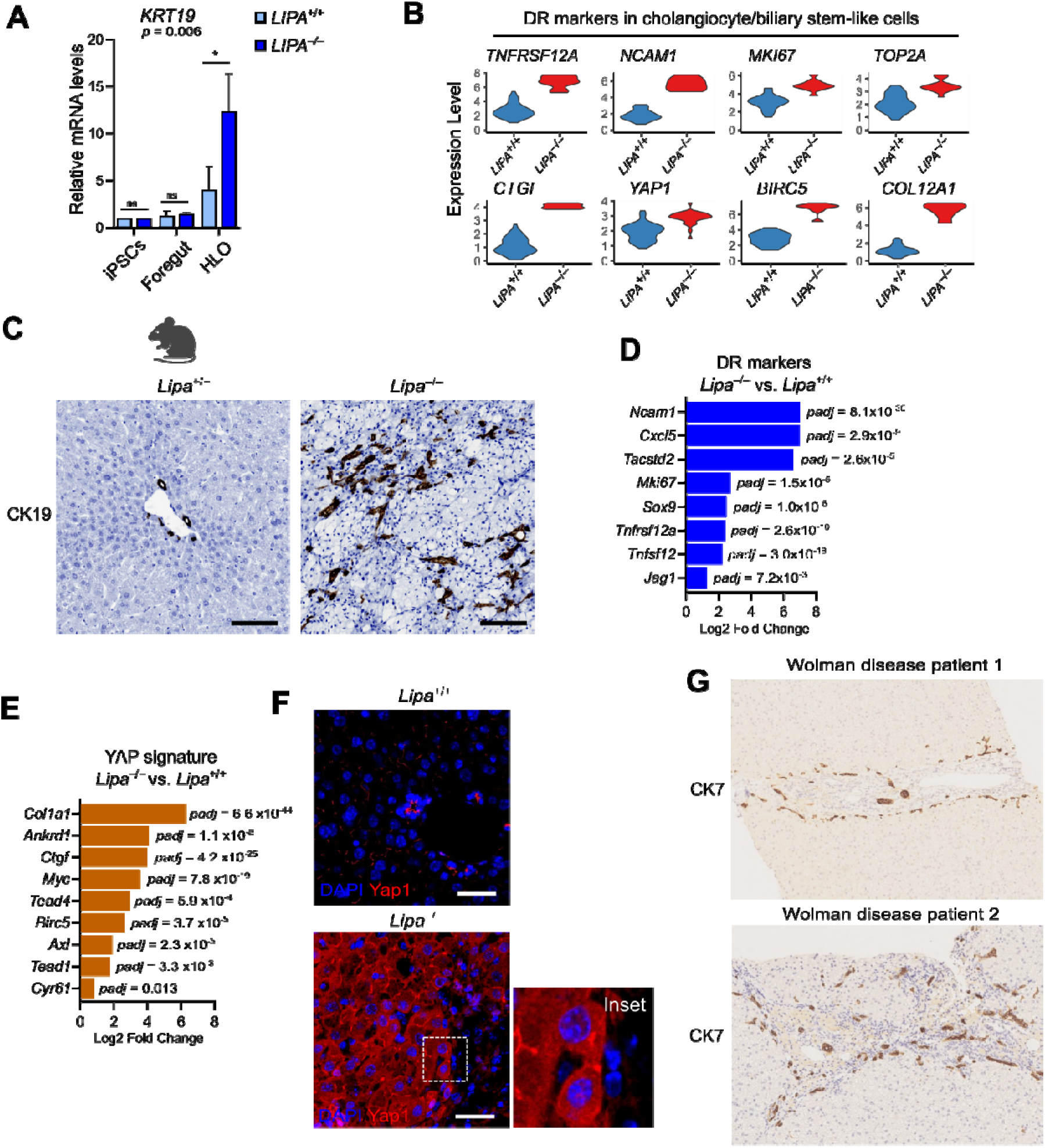
HLO capture ductular reactive states emerging in WD. **A** Quantitative RT–PCR analysis of *KRT19* in *LIPA^+/+^* and *LIPA^-/-^* cells at day 0 (iPSCs), 7 (foregut) and 21 (HLO). Results represent mean ± SD; n = 4-6 independent differentiation experiments. Two-way ANOVA was applied followed by Tukey post-hoc test. ns = not significant, * *p*□≤□0.05. **B** Violin plots showing expression of ductular reaction (DR) markers in cholangiocyte-like and biliary tree stem-like cells grouped by genotype. **C** Representative IHC staining of CK19 in livers from adult *Lipa^+/+^*and *Lipa^-/-^*mice. The scale bar represents 100 μm. **D** Upregulated DR markers in whole liver of WD mice from Laurent et al. **E** Upregulated Yap1 target genes in whole liver of WD mice from Laurent et al. **F** Representative IF staining of YAP1 in livers from Lipa^+/+^ and Lipa^-/-^ mice. The scale bar represents 50 μm. **G** Representative IHC staining of CK7 in livers from Wolman disease patients.

To determine whether the reactive biliary phenotype captured by WD organoids reflected an authentic tissue-level manifestation of WD, rather than an organoid-specific *in vitro* response, we examined liver tissue from WD mice^7^. Remarkably, Krt19 and Hnf4α immunostaining confirmed a pronounced expansion of the biliary compartment and concomitant loss of hepatocyte identity respectively (Figures 5C and Supplementary Figure 5B, C). Consistently, bulk RNA sequencing of *Lipa^-/-^* livers revealed a robust upregulation of DR-associated genes, mirroring the transcriptional changes identified in HLO (Figure 5D)^7^. Moreover, livers from WD mice showed upregulation of the YAP signature and enhanced YAP protein activation (Figure 5E, F).

Finally, to assess whether the reactive biliary phenotype captured in WD organoids reflects a *bona fide* histological feature of human disease, we examined liver tissue from WD patients by immunostaining for the biliary marker KRT7. Strikingly, portal regions displayed marked expansion and remodeling of the biliary epithelium, with enrichment of irregular neo-ductular structures characterized by small or inconspicuous lumina and localization along the limiting plate (Figures 5G and Supplementary Figure 5D). These histological features were consistent with a reactive ductular response and mirrored the alterations observed in both organoids and mice.

Together, these observations support the ability of multicellular HLOs to reproduce complex tissue-level biliary responses, such as the DR, emerging in the WD liver microenvironment.

### Hepatocyte-directed LIPA restoration suppresses reactive biliary responses *in vivo*

To clarify whether the DR-like program observed in WD organoids was driven by cell-autonomous LIPA deficiency in the biliary lineage—affecting either cholangiocyte development or mature cholangiocyte behaviour—or instead represented a non-cell-autonomous adaptive response of the biliary compartment to parenchymal injury, we first assessed bile duct development in WD mice. Interestingly, we found no major abnormalities in biliary development in foetal as well as in post-natal livers (Supplementary Figure 5F). Next, we investigated whether LIPA loss directly affects adult biliary epithelial behaviour in a cell-autonomous manner. To this aim, we derived biliary organoids from wild-type and WD mice and cultured them over multiple passages as cholangiocyte-only epithelial cultures, thereby isolating biliary-intrinsic responses from the influence of hepatocyte and stromal compartments (Figure 6A). Despite the complete lack of *LIPA* expression (Supplementary Figure 6A), we could not identify any proliferation, viability or morphological abnormality in WD organoids (Figure 6B, C, E). Accordingly, markers of DR were not activated in WD organoids (Figure 6D). To corroborate these data in human context, we next generated human liver–derived biliary organoids and engineered them by CRISPR/Cas9 to disrupt the LIPA gene (Supplementary Figure 6B, C)^9,45^. Efficient reduction of LIPA protein expression was confirmed, and LAMP2 staining demonstrated lysosomal accumulation (Supplementary Figure 6D, E). However, LIPA-deficient organoids did not show induction of DR-associated genes (Supplementary Figure 6F), nor evident changes in morphology or proliferative behaviour (Figure 6F), suggesting that the DR observed in HLO and *in vivo* depends on non-cell-autonomous tissue response to injury^39,40^. To directly test this possibility in an *in vivo* gene therapy setting, we analyzed liver samples from hepatocyte-targeted AAV-LIPA–treated *Lipa^-/-^* mice, in which wild-type human LIPA expression had been restored in the hepatic parenchyma in previous experiments^7^. Consistent with hepatocyte-directed transgene expression, HA-tagged LIPA was detected in hepatocytes but not in Krt19□ cholangiocytes, as shown by combined HA and Krt19 staining (Figure 6G). Notably, hepatocyte-restricted restoration of LIPA was accompanied by a substantial suppression of DR-associated markers compared with untreated mice (Supplementary Figure 6G, H). Histological analysis further revealed a marked reduction in Krt19□ ductular expansion and YAP activation, indicating attenuation of the reactive biliary response following correction of hepatocellular LIPA deficiency (Figures 6G and Supplementary Figure 6I).

**Fig. 6:**
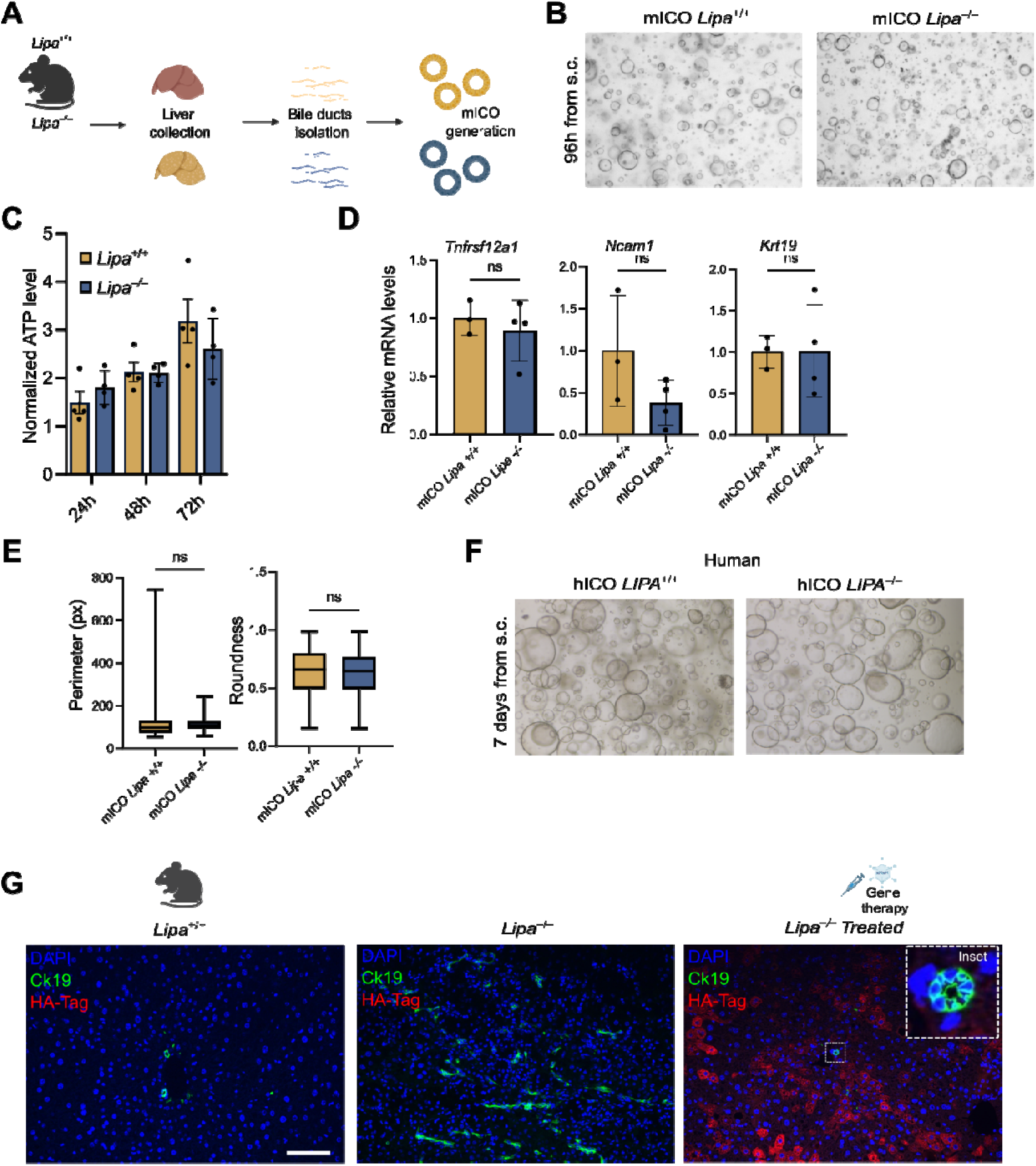
**Hepatocyte-directed LIPA restoration suppresses reactive biliary responses in vivo**. A Schematic representation of the mouse cholangiocyte organoids (mICO) generation from *Lipa^+/+^* and *Lipa^-/-^* mice. **B** Bright-field image of BME drop containing *Lipa^+/+^* and *Lipa^-/-^* mICO. **C** ATP-based viability assay in *Lipa^+/+^* and *Lipa^-/-^* at different timepoints of organoid growth. Results represent mean ± SEM; n = 4. Two-way ANOVA: time, p = 0.009; genotype, ns; time × genotype, ns. Tukey’s post-hoc test: ns. ns = not significant. **D** Quantitative RT–PCR analysis of *Lipa* and DR markers in *Lipa^+/+^*and *Lipa^-/-^* mICO. Results represent mean ± SD; n = 3. Student’s t-test was applied: ns = not significant. **E** Quantification of shape/morphological descriptors in *Lipa^+/+^* and *Lipa^-/-^* mICO at 96 hours after single cell dissociation. Results represent mean ± SD. Student’s t-test was applied: ns = not significant. **F** Bright-field image of BME drop containing *LIPA^+/+^* and *LIPA^-/-^*hICO. **G** Representative IF staining of Krt19 and HA-Tag in liver section of *Lipa^+/+^, Lipa^-/-^* and *Lipa^-/-^* AAV treated mice. The scale bar represents 100 μm.

Together, these findings indicate that DR in WD is primarily driven by hepatocellular injury through non-cell-autonomous tissue responses, rather than by an isolated biliary cell-autonomous defect. The ability of multicellular HLO to recapitulate this emergent reactive biliary program underscores their capacity to model the crosstalk between parenchymal damage, biliary remodeling, and injury-associated tissue dynamics.

### AAV-mediated *LIPA* restoration in hepatocyte-like cells attenuates disease responses in WD organoids

Having established that reactive biliary responses arise as a consequence of hepatocellular LIPA deficiency and can be suppressed by hepatocyte-directed gene therapy in vivo, we next asked whether HLO could also recapitulate this therapeutic response. If HLO faithfully model the complex cellular interactions underlying Wolman disease, restoration of LIPA selectively in hepatocyte-like cells should be sufficient to remodel not only hepatocellular defects but also the higher-order pathological programs emerging within the organoid microenvironment, including inflammatory, fibrogenic and reactive biliary responses. To explore this possibility, we first evaluated the transduction efficiency of multiple AAV serotypes in HLO. Organoids were transduced with a panel of GFP-expressing AAV vectors (Figure 7A), among which AAV6 consistently produced the strongest GFP signal by fluorescence microscopy (Figure 7B), in agreement with previous reports in human liver organoid systems^46^. We therefore selected AAV6 as an enabling vector for therapeutic gene delivery in WD organoids. To assess whether parenchymal restoration of LIPA activity could reverse multicellular disease-associated programs, we transduced LIPA^-/-^ HLOs with an AAV6 vector carrying wild-type human LIPA under the control of the liver-associated *SERPINA1* (*AAT1*) promoter (Figure 7C), which is predominantly active in hepatocyte-like cells within HLOs (Supplementary Figure 7A). Successful transgene delivery was confirmed by robust restoration of LIPA expression (Supplementary Figures 7B, C). Importantly, AAV-mediated LIPA restoration significantly attenuated molecular signatures associated with metabolic stress, inflammation, fibrosis and DR in WD organoids (Figure 7D, E). These findings were consistently observed across independent HLO differentiation experiments. Collectively, these findings indicate that multicellular disease states emerging in WD organoids can be therapeutically modulated and establish HLO as a human platform to evaluate tissue-level therapeutic rescue in a multicellular hepatic context.

**Fig. 7:**
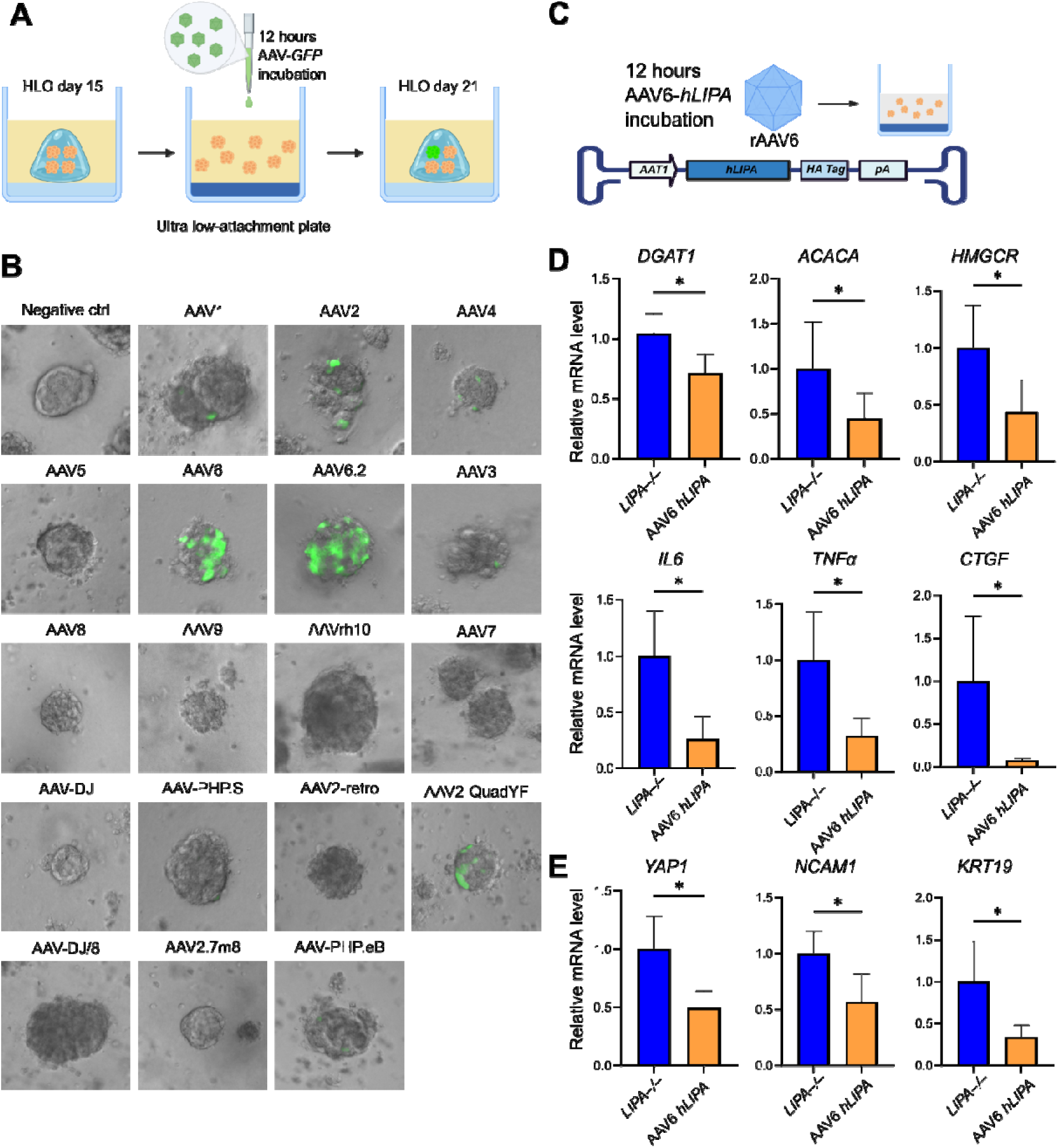
HLO provide a preclinical platform to evaluate AAV-mediated gene therapy in Wolman disease. **A** Schematic representation of the experimental workflow for AAVs infection in HLOs. **B** Bright-field and fluorescence images of HLOs infected with the 18 AAV serotypes expressing GFP (green). **C** Schematic representation of the recombinant AAV6-*hLIPA* vector design and infection protocol. **D** Quantitative RT–PCR analysis of lipid metabolism, fibrosis and inflammation in *LIPA^-/-^*and AAV6-*hLIPA* HLOs. Results represent mean ± SD; n = 4-6 independent differentiation experiments. Student’s t-test was applied: * *p*□≤□0.05. **E** Quantitative RT–PCR analysis of DR in *LIPA^-/-^* and AAV6-*hLIPA* HLOs. Results represent mean ± SD; n = 4-6 independent differentiation experiments. Student’s t-test was applied: * *p*□≤□0.05.

## Discussion

The human liver organoid model of WD described here provides a significant advance over existing models. By using CRISPR/Cas9 to generate an isogenic *LIPA* knockout in a well-defined hiPSC line, we isolated the impact of *LIPA* deficiency on a uniform genetic background, avoiding the variability inherent to patient-derived cells.

The resulting hepatic organoids recapitulate key biochemical, cellular, and transcriptional features of WD, including loss of LAL activity, neutral lipid accumulation, lysosomal expansion, metabolic stress, and activation of inflammatory and fibrogenic programs associated with WD liver pathology. Their multicellular organization, comprising hepatocyte-like parenchymal cells, cholangiocytes, and stromal cells, enabled the identification of cell type–specific disease-associated states that are difficult to capture in simplified monoculture systems. In hepatocyte-like cells, LIPA deficiency induced profound metabolic rewiring, whereas stromal populations acquired transcriptional signatures associated with extracellular matrix remodelling, TGFβ signalling, inflammation, and fibrogenic activation, closely mirroring processes observed in WD liver tissue.

Of note, biliary cells in WD organoids acquired a reactive transcriptional state consistent with DR, a tissue response commonly associated with chronic liver injury and regenerative remodelling. Examination of liver tissue from WD patients revealed expansion and remodelling of KRT7□ ductular structures closely mirroring the alterations observea in HLO. Together, these findings support the ability of HLO to reproduce reactive biliary responses emerging in the diseased hepatic microenvironment.

To investigate the contribution of epithelial compartmental crosstalk to this phenotype, we generated human *LIPA*^-/-^ biliary organoids. To our knowledge, this represents the first direct assessment of the cell-specific consequences of LIPA loss in human biliary cells. Although these organoids exhibited evidence of lysosomal dysfunction, they failed to induce DR-associated transcriptional programs or clear proliferative changes. These findings suggest that LIPA loss in biliary cells alone is insufficient to efficiently reproduce the reactive biliary state observed in HLO. Instead, they support the notion that DR in WD emerges in close association with intercellular communication and parenchymal stress signals within the hepatic microenvironment. This capacity of HLOs to recapitulate emergent and physiologically relevant tissue interactions represents a major strength of the system.

Additional support for this model emerged from our *in vivo* studies in *Lipa*^-/-^ mice and patients tissues, which showed a DR closely resembling the phenotype observed in WD organoids and patient tissues. Importantly, hepatocyte-directed AAV-mediated restoration of human *LIPA* strongly suppressed DR-associated transcriptional programs and Krt19□ ductular expansion. This genetically defined rescue strategy provides a uniquely clean experimental framework to investigate the origin of DR, a process widely proposed to arise as a secondary response to parenchymal injury but not yet formally demonstrated, largely because reactive biliary responses are typically studied in complex multifactorial liver diseases that affect multiple hepatic compartments simultaneously. In such contexts, disentangling whether DR arises from biliary-intrinsic injury or instead represents a secondary response to parenchymal dysfunction is inherently challenging. Here, selective correction of LIPA deficiency in hepatocytes was sufficient to attenuate the biliary reactive response, confirming a model in which cholangiocytes sense and respond plastically to parenchymal injury rather than undergoing DR as a direct consequence of biliary-intrinsic damage.

Beyond modelling WD pathology, this platform also provides an opportunity to evaluate therapeutic modulation of disease-associated programs in a human hepatic-like context. Leveraging HLO as a preclinical testbed, we evaluated the transduction efficiency of multiple AAV serotypes and identified AAV6 as an efficient vector for gene delivery into WD organoids. Restoration of LIPA expression by AAV-mediated gene transfer attenuated transcriptional programs associated with metabolic stress, inflammation, fibrosis, and reactive biliary remodelling. These findings indicate that disease states emerging in WD organoids can be therapeutically modulated, extending the utility of this platform beyond modelling toward evaluation of tissue-level rescue.

Given that the liver represents both a major target and a major source of toxicity in systemic AAV-based therapies, liver organoids may additionally provide a human platform to evaluate vector tropism and potential hepatotoxicity in a physiologically relevant context^47^. Moreover, beyond AAV vectors, WD organoids could be exploited to assess alternative delivery systems, including lentiviral vectors and lipid nanoparticle–based approaches. Finally, the combination of isogenic engineering and tissue-like organization makes these organoids particularly well suited for CRISPR-based functional genomics and pharmacological screening approaches aimed at identifying pathways that regulate disease progression or repair.

Despite these advantages, the model still has intrinsic limitations that could be further optimized. While HLOs reproduce the major hepatic lineages, they lack a fully developed vascular compartment and mature endothelial cells, which are important contributors to liver homeostasis, immune crosstalk, and vector biodistribution. In addition, immune populations are not represented in this system, limiting its ability to capture inflammatory and immune-mediated interactions that contribute to liver pathology. Likewise, the relative abundance of parenchymal, biliary, and stromal populations does not fully reflect the composition of native human liver tissue, potentially influencing tissue-level responses. Nonetheless, the convergence between organoid (bulk and single-cell analysis), mouse, and patient data supports the biological relevance of the reactive biliary responses identified in this study.

In sum, this study highlights the ability of human liver organoids to model emergent tissue-level responses associated with WD, including reactive biliary remodeling arising from dynamic hepatic compartmental crosstalk. Beyond recapitulating key metabolic and inflammatory features of disease, this platform enables the investigation and therapeutic modulation of higher-order pathological programs within a human tissue-like context. These findings support the broader utility of organoid systems for studying complex liver diseases and for evaluating tissue-level therapeutic rescue strategies.

## Material & methods

### Human iPSCs culture and maintenance

The *LIPA^+/+^* human induced pluripotent stem cell (hiPSC) line used in this study was obtained from ThermoFisher Scientific (Gibco, A18945; Monza, Italy) and verified for pluripotency and normal karyotype. This hiPSC line was edited by CRISPR/Cas9 editing (see below) to generate *LIPA^-/-^*hiPSCs. Both hiPSC lines were maintained under feeder-free conditions in Essential 8 Medium (Thermo Fisher Scientific, A1517001) on 6-well plates coated with Culturex Basement Membrane Extract (BME) Type 2 (Biotechne, 3533-010-02) diluted 1:60 in Adv/DMEM-F12 medium (Sigma-Aldrich, SCM162). Cultures were incubated at 37 °C in a humidified atmosphere of 5% CO□ and 95% air, and the medium was replaced three times a week. For passaging, cells at ∼70–80% confluence were washed twice with PBS (1 mL/well) at room temperature and incubated with 0.5 mL of Versene (Gibco, 15040-066) for 2-3 min at 37 °C. The Versene solution was aspirated, and fresh Essential 8 Medium was added. Colonies were detached by gentle pipetting to generate small aggregates, and an appropriate volume of the suspension was transferred to a new BME-coated plate. To enhance post-dissociation survival, the medium was supplemented with 10 µM Y-27632 (MCE, HY-10583) for the first 24 h after plating. The following day, the medium was replaced with fresh Essential 8 Medium lacking Y-27632. To generate the different clonal lines, hiPSCs were dissociated into a single-cell suspension and plated at low density to enable the isolation of cells, which were then expanded and genotyped by Sanger sequencing.

### Foregut induction and generation of human liver organoids (HLO)

*LIPA^+/+^* and *LIPA^-/-^* hiPSCs were maintained under feeder-free conditions as described above and differentiated into foregut following a previously published protocol with minimal^10–12^. In brief, hiPSCs were dissociated using Versene (Gibco, 15040-066) and seeded onto BME coated plates at a density of 7.5×10□ cells/well in Essential 8 medium supplemented with 10 µM Y-27632 (MCE, HY-10583) for the first 24h. Differentiation was initiated by replacing the medium with RPMI 1640 (Thermo Fisher Scientific, 11875093) containing 100 ng/mL Activin A (Proteintech, HZ-1138) and 50 ng/mL bone morphogenetic protein 4 (BMP4; Proteintech, HZ-1045) on day 1. On day 2, cells were cultured in RPMI 1640 supplemented with 100 ng/mL Activin A and 0.2% fetal bovine serum (FBS; Euroclone, ECS0180L), followed by 100 ng/mL Activin A and 2% FBS on day 3. From days 4 to 6, cells were cultured in Adv/DMEM-F12 medium (Sigma-Aldrich, SCM162) supplemented with B27 (Thermo Fisher Scientific, 12587010), N2 (Thermo Fisher Scientific, 17502048), 500 ng/mL fibroblast growth factor 4 (FGF4; GenScript, z02984), and 3 µM CHIR99021 (BioGems, 2520691). Cultures were maintained at 37 °C in 5% CO□ with daily medium replacement. At day 7, foregut progenitors were detached and dissociated using TriplE Express (Gibco, 12604013) and cryopreserved in FBS with 10% DMSO at −80 °C or liquid nitrogen for long-term storage. For liver organoid generation, frozen foregut cells were rapidly thawed, centrifuged at 180 x g for 3 min, and 1×10^6^ cells were resuspended in a dome composed of 50% BME and 50% “five factors” medium composed as follow: Adv/DMEM-F12 (Sigma-Aldrich, SCM162) supplemented with 2% B27 (Thermo Fisher Scientific, 12587010), 1% N2 (Thermo Fisher Scientific, 17502048), 10 mM HEPES (Sigma-Aldrich, 83264), 1% L-Glutamine (Euroclone, ECB3004D), 1% Penicillin–Streptomycin (Sigma-Aldrich, P4333), 5 ng/mL fibroblast growth factor 2 (FGF2; ACRO biosystems, BFF-H4117), 10 ng/mL vascular endothelial growth factor (VEGF; Proteintech HZ-1038), 20 ng/mL epidermal growth factor (EGF; Peprotech, 315-09), 3 µM CHIR99021 (BioGems, 2520691), 0.5 µM A83-01 (BioGems, 9094360), and 50 µg/mL ascorbic acid (Sigma-Aldrich). Cultures were maintained at 37 °C in 5% CO□ in “five factors” medium with medium changes every 2 days for 4 days. The medium was then replaced with Adv/DMEM-F12 with 2% B27, 1% N2, 10 mM HEPES, 1% L-Glutamine, 1% Pen/Strep, and 2 μM Retinoic Acid (RA; Sigma-Aldrich, R2625), and incubated in the CO_2_ incubator for further 4 days with medium changed every 2 days. After 4 days the organoids were gently collected using cell recovery solution (Corning, 354253), filtered through a 150-µM cell strainer and replated with Hepatocyte Culture Medium (HCM; Lonza, MD, USA, CC-3198). The cells were incubated in a CO_2_ incubator for 6 days, changing the medium every 2 days. At day 21, mature HLO were collected and used for subsequent experiments.

### Human and mouse liver samples

Samples for hICO and histological analysis were generated from a patient undergoing bariatric surgery diagnosed with a mild steatosis (Kleiner score = 1). Sensitive data were protected through anonymization. Diagnostic CK7 immunoistochemistry of Wolman disease patients were obtained from the Department of Pathology, Hôpital Universitaire Necker-Enfants Malades, APHP, Paris, France.

Heterozygous Lipa^tm1a(EUCOMM)Hmgu/Biat^ mice were crossed with heterozygous mice expressing Cre recombinase under the control of the *CMV* promoter to delete exon 4 of the murine *LIPA* gene. Homozygous mice were subsequently backcrossed onto a C57BL/6N background for more than eight generations. Animals were maintained under specific pathogen-free (SPF) conditions with a 12-hour light/dark cycle and ad libitum access to standard chow and water. All animal experiments were performed in accordance with French and European regulations on animal experimentation. Fresh liver samples from wild-type (*Lipa^+/+^*) and *Lipa*^□/□^ mice were collected at 22±2 and 44±2 weeks. A total of 40 mg of frozen liver tissue was sent to Azenta Life Science (Leipzig, Germany) for RNA sequencing, which was performed as previously described^7^.

### Human and mouse cholangiocyte organoids generation

Intraepatic cholangiocyte organoids (iCOs) were isolated from either *Lipa^+/+^* and *Lipa*^□/□^ mice or human biopsies following established protocols^45^ with minor modifications. Briefly, mouse livers were stored in Advanced DMEM/F12 supplemented with 1% pen/strep, 1% L-glutamine, and 1% HEPES at 4□°C for 24□h prior to processing to allow transport from France to Italy. Bile ducts were then extracted by incubating the minced liver pieces in digestion buffer (0.125 mg/mL collagenase and 0.125 mg/mL dispase II in DMEM) for 45–120 min at 37°C under continuous agitation. Ductal structures were separated from the undigested tissue by centrifugation and resuspended 25 μL BME droplets in 24-well plates. Human biopsies were placed in William’s E medium and transported to the laboratory on ice. Liver samples were then minced into small-cell clusters and digested in human digestion buffer (2.5 mg/ml collagenase D (Sigma-Aldrich, 11088866001) and 0.1 mg/ml DNaseI in Hanks’ balanced salt solution) at 37□°C for 75-120 minutes under continuous agitation. Cell clusters were separated from the undigested tissue by centrifugation and resuspended 25 μl BME droplets in 24-well plates. Following BME solidification, 500 μl of either mouse or human liver isolation media was added. Once the bile ducts started budding, isolation medium was substituted with liver expansion medium. The medium was changed every 2–3 days. Fully grown organoids were enzymatically dissociated as single cells using TrypLE Express (Gibco, 12604013) before being resuspended in BME and cultured in respective expansion medium supplemented with 10 μM Y-27632 (MCE, HY-10583) for the initial 3 days.

### LIPA gene editing with CRISPR/Cas9

*LIPA*□^/^□ cells were generated by exploiting the Lonza 4D-Nucleofector® System (Lonza Bioscience) according to manufacturer’s instruction and adapting the protocol described in Ghetti et al, 2021 to iPSCs^20^. A gRNA targeting exon 4 of LIPA gene (5’GCTGGCAGATTCTAGTAACT3’) was obtained annealing 1:1 100 µM Alt-R® CRISPR-Cas9 crRNA (IDT™) to 100 µM Alt-R® CRISPR-Cas9 tracrRNA (IDT™ Cat# 1072532). Ribonucleoparticles (RNPs) were prepared mixing 150 pmol of gRNA and 120 pmol of recombinant Cas9. iPSCs were pre-treated with 10 µM Y-27632 (MCE, HY-10583) for 30 minutes, then dissociated to single cells with TrypLE™ Express Enzyme (Gibco™ 12605010); 2.5×10□ cells/reaction were resuspended in P3 Primary Cell Full Electroporation Buffer (Lonza Bioscience, kit V4XP-3032) and added to a mix containing the RNP and 4 µM Alt-R® Cas9 Electroporation Enhancer (IDT™ Cat# 1075915). Cells were electroporated using Lonza 4D-Nucleofector CM-113 protocol and then plated in presence of 10 µM Y-27632. After 24 hours Y-27632 concentration was halved: the drug was completely removed the day after.

The genomic DNA was extracted from iPSCs and hICOs using NucleoSpin® Tissue kit (Macherey-Nagel 740952) following manufacturer protocol. PCR products spanning the edited site were obtained using PCR primer pair FW (5’TGGTCCAGTTTCACTGCTACCTT3’) – RV (5’AATAACGTTTACCCAGCAGCAC3’), Sanger sequenced and analyzed with ICE online tool (Synthego).

### LIPA activity measurements

Human iPSCs were homogenized in PBS containing 0.5% Triton X-100 and a protease inhibitor cocktail (Roche), followed by centrifugation at 13,000 × *g* for 10 min at 4 °C. Protein concentration in the supernatant was determined using the BCA Protein Assay Kit (Thermo Fisher Scientific). For lysosomal acid lipase (LAL) activity measurements, 7µg of total protein was used per reaction. Enzymatic activity was assessed as previously described^7^. Briefly, samples were incubated for 10 min at 37 °C with either 42 µM Lalistat-2 (Sigma-Aldrich, St. Louis, MO, USA) or water and then transferred to an OptiPlate-96 F plate (PerkinElmer). Reactions were initiated by adding 75 µL of substrate buffer containing 340 µM 4-MUP, 0.9% Triton X-100, and 220 µM cardiolipin in 135 mM acetate buffer (pH 4.0). Fluorescence was recorded at 37°C over 35 cycles at 30-second intervals using a SPARK TECAN Reader (Tecan). Kinetic parameters (average rate) were calculated with Magellan Software, and LAL activity was quantified according to Equation 1.

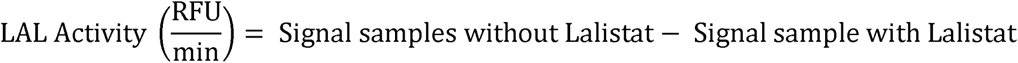

### Flow Cytometry

hiPSCs were dissociated to single cells by the treatment of Versene for 5 min. After PBS wash, the single cells were fixed with 4% PFA and washed 3 times in PBS. Cells were incubated for 30 minutes with Nile Red 1:10,000 in PBS and subjected to flow cytometry. Data were acquired on FACSCelesta (BD Biosciences) and analyzed using FlowJo software (FlowJo, Ashland, OR, USA).

### Histological analysis

HLO domes were fixed in 2% PFA with 0.1% glutaraldehyde for 20 minutes at room temperature and washed 2 times with PBS at room temperature. Following incubation with an increasing series of ethanol and xylene, HLOs domes were embedded in paraffin. 5-µm-thick organoid sections were processed for hematoxylin-eosin (H&E) and Sirius red staining.

Fresh liver of 44 ± 2-week-old WT*, Lipa*□^/^□ and *Lipa*□^/^□ Treated mice were fixed in formaldehyde and subsequently embedded in paraffin. Serial 4μm cross-sections were prepared using a Leica RM2245 microtome (Leica Biosystems). To reduce sampling bias, two sections were collected from each specimen. Similarly, the human sample was fixed in 10% formalin for 24 hours at RT. Following incubation with an increasing series of ethanol and xylene, the sample was embedded in paraffin.

### Immunohistochemistry

Immunohistochemical staining was performed on 4-µm-thick mouse liver sections. After dewaxing and rehydration, slides underwent antigen retrieval in heated 10 mM sodium citrate tribasic, 0.05% Tween 20 solution (pH 6.0) or ImmunoDNA Retriever 20X with EDTA (BSB 0030) for 30 minutes. Endogenous peroxidase activity was blocked with 3% hydrogen peroxide solution for 10 minutes, and unspecific bounds with 1% BSA, 0.01% Triton X-100 in PBS for 1 hour at RT. Samples were incubated with Ck19 or CK7 primary antibody diluted 1:100 overnight in a humidified chamber at 4°C. The following day, slides were washed with PBS and incubated with HRP-conjugated secondary antibody (Life Technology) diluted 1:200 for 2 hours. After two washes with PBS, samples were incubated with a DAB peroxidase substrate kit (Vector Laboratories, SK-4100) and counterstain with Mayer’s hematoxylin (Sigma, MHS1). All antibodies used in this article are listed in Supplementary Table 1.

### Immunohistochemistry quantification

For the quantification of Ck19+ mouse liver areas, stained tissue samples were scanned with an Axioscan 7 (Zeiss, Germany). Ck19+ area was calculated using QuPath software (version 0.5.1) and represented as a percentage of the total hepatic tissue area. The graphed data shows the mean ± SD from 4 independent experiments.

### Immunofluorescence

For HLOs the immunostaining was performed on 5-µm-thick organoid sections obtained from paraffin-embedded blocks. De-paraffinization and antigen retrieval were carried out as described in the previous paragraph. After blocking unspecific bindings with 1% BSA and 0.01% Triton X-100 in PBS for 1 hour at RT, tissues were incubated with primary antibodies (1:100 dilution) overnight in a humidified chamber at 4°C. The following day, slides were washed with PBS and incubated with specific secondary antibodies conjugated with Alexa Fluor 488 or 594 (Jackson Immuno Research; 1:200) in blocking solution for 2 hours at RT. Nuclei were stained with 1 µg/mL DAPI, and ProLong Gold antifade reagent (Life Technology, P36934) was used to mount coverslips. For 2D immunofluorescence on hiPSCs, cells were seeded on BME-coated coverslips and fixed at ∼80% confluency with 4% paraformaldehyde for 20 minutes at RT. After three washes in PBS, cells were permeabilized with 0.2% Triton X-100 in PBS for 1 hour, followed by blocking with 1% BSA and 0.01% Triton X-100 for 1 hour at RT. Primary antibodies (SOX2, LAMP2, YAP1, HA-Tag, KRT19 and HNF4A, 1:100) were applied overnight at 4°C in a humidified chamber, followed by incubation with secondary antibodies (1:500) for 2 hours at RT. Nuclei were counterstained with DAPI prior to mounting. For LIPA immunofluorescence, hiPSC-derived cells grown on coverslips were fixed in cold methanol, followed by blocking, and antibody incubation steps performed as described above. Representative images shown are from experiments independently repeated at least three times with consistent results.

### Image acquisition and organoids shape quantification

Immunofluorescence images were acquired at confocal microscope (Zeiss, LSM 880 Airyscan) or with a ZEISS Apotome 3. Bright-field images of hiPSCs, HLOs, mICOs and hICOs as well as GFP positive HLO images were acquired using EVOS™ M5000 Imaging System (AMF5000). Images were processed using ImageJ software (version 1.54f). The analysis of organoid morphology descriptors was performed using ImageJ software (version 1.54f), with the BetterWand function employed to analyze and quantify organoid roundness, perimeter and area.

### Viability assay

Cell viability was assessed at 3 different timepoints (24h, 48h and 72h) in *Lipa*^+/+^ and *Lipa*^-/-^ mICO by measuring ATP levels with the CellTiter-Glo luminescent cell viability kit (Promega, G6081) following the manufacturer’s instructions. Data are reported as normalized ATP level in 4 biological replicates.

### Quantitative real-time qRT-PCR for mRNA quantification

RNA was extracted from organoids using the NucleSpin Tissue kit (Macherey-Nagel, 740952) following the manufacturer’s instructions. Reverse transcription of total RNA was performed with the high-capacity RNA-to-cDNA kit (Applied biosystem, 4387406) following manufacturer’s instructions. The expression of selected genes was analyzed using the CFX Duet real-time PCR detection system (Bio-Rad, USA) and the PowerTrack SYBR green master mix (Applied biosystem, A46109). Expression values were calculated using the ΔΔCt method with the CFX Maestro 2.2 Manager software (Bio-Rad, USA), and gene expression levels were normalized against ACTB and GAPDH as the housekeeping genes. Primers for qRT-PCR are listed in Supplementary Table 2.

### Western blotting

Total proteins were isolated from organoids using lysis buffer (50 mM Tris-HCl (pH 8), 150 mM NaCl, 1 mM EDTA, 0.5% Nonidet P-40, 0.05% sodium deoxycholate, and protease and phosphatase inhibitors) followed by sonication and centrifugation for 10 min at 12 000g at 4°C. Protein concentration was determined using the BCA Protein Kit Assay (Sigma-Aldrich, 71285-3), according to the manufacturer’s instructions. A total of 20 µg of proteins was separated by SDS–PAGE and transferred onto nitrocellulose membranes (Millipore). After blocking with 5% milk in TBST for 1 hour at RT, membranes were incubated overnight with primary antibodies diluted in blocking buffer. Anti-LIPA antibody was used at 1:200 for hICOs and 1:500 for hiPSCs, while HRP-conjugated Vinculin was used as loading control (1:2 000). The following day, membranes were incubated with HRP-conjugated secondary antibodies (Life Technology) at 1:1 000–1:2 000 for 1 hour at RT. Protein detection was performed using the ECL kit (Thermo Fisher Scientific, 32132) and visualized on a ChemiDoc MP Imaging System (Bio-Rad, USA).

### Single cell RNA sequencing library preparation

The scRNA-seq libraries were constructed using the GEXSCOPE® Single Cell RNA Library Kit (Singleron Biotechnology) following the manufacturer’s protocol for a minimal capture of 10.000 cells. For each genotype, a single scRNA-seq sample was generated by pooling multiple organoids and was subsequently subjected to scRNA-seq analysis. Illumina-compatible NGS libraries generated were sequenced on an Illumina NovaSeq 6000 instrument using a paired-end 150bp approach. The reads were demultiplexed on Illumina’s BaseCloud, and fastq files were used to initiate data analysis.

### Data processing and QC

All the original samples generated for this paper were mapped and expression levels summarized using Celescope pipeline version 2.3.0. Human samples were mapped against the GRCh38.v1.8.0 version of the Homo sapiens genome, employing default parameters. A total of 9041 *LIPA^+/+^* HLO and 4331 *LIPA^-/-^* HLO cells were identified. Seurat (v5.2.1) objects were created using genes expressed in more than three cells and cells with more than 200 features expressed. Cells with fewer than 750 features, fewer than 1,500 counts, or more than 20% of counts mapping to mitochondrial genes were excluded. The number of *LIPA^-/-^* cells retained after quality control was lower than that of wild-type cells, likely reflecting reduced recovery of viable mutant cells during single-cell suspension preparation and library generation. This may be due to the increased sensitivity of metabolically stressed LIPA-deficient cells to the mechanical and enzymatic stress imposed by tissue dissociation. Nevertheless, the dataset retained sufficient depth for exploratory single-cell analyses, and the main biological findings were subsequently further validated using independent molecular and cellular assays across multiple experimental models.

### Normalization and annotation of cells

Data was normalized using Seurat SCTransform function. Principal Component Analysis (PCA) was performed using the top 2,000 highly variable genes, and the top 20 principal components (PCs) were selected based on an elbow plot of explained variance. Clustering was performed using Seurat’s FindNeighbors and FindClusters functions, with parameters selected according to the dataset. The clustering results were visualized using UMAP dimensionality reduction. Differential expression analysis was performed using Seurat’s FindAllMarkers function, with genes exhibiting adjusted p-values < 0.05 considered significant. Marker genes for each cluster were identified and analyzed to uncover cluster-specific signatures. Detailed marker lists per cluster can be found in Supplementary Table 3.

### Publicly available dataset analysis

The human scRNA-seq dataset from Ouchi et al. (GSE130073) and our *LIPA^+/+^* scRNA-seq dataset were merged into a single Seurat object. The merged datasets were normalized using the SCTransform function, and dimensionality reduction was first performed with principal component analysis (PCA; 50 principal components). To mitigate potential batch effects across datasets, we applied the RunHarmony function. Finally, UMAP dimensionality reduction was used for visualization of the harmonized data. Processed bulk RNA-seq data from 8-month-old WT*, Lipa*□/□ and *Lipa*□/□ Treated mice were downloaded from GEO (GSE252742).

### Lipid Extraction

Lipids were extracted from cell pellets using a chloroform/methanol/water protocol. Cells pellets were resuspended with 100 µL of methanol containing 5% SPLASH internal standard, followed by the addition of 750 µL of chloroform. Samples were sonicated for 2 min in an ultrasonic bath and mixed for 6 min at 2000 rpm in a thermoblock at 4 °C. Then, 100 µL of water were added, and the mixture was centrifuged at 14,000 rpm for 2 min to induce phase separation. The lower organic phase (500 µL) was collected, transferred to a clean microtube, and dried in a speedvacum system. The dried extract was stored at-80°C and resuspended in 100 µL of methanol + CUDA an internal standard, before the LC-MS analysis.

### Lipidomic Analysis

Lipidomic analysis was performed with an UHPLC Vanquish system (Thermo Scientific, Rodano, Italy) coupled with an Orbitrap Q-Exactive Plus (Thermo Scientific, Rodano, Italy). An Acquity UPLC CSH C18 column (1.7 µm, 2.1 × 150 mm) was used, with the column temperature maintained at 40 °C, and a constant flow rate of 0.200 mL min□¹. Mobile phase A was H□O/MeOH (50:50, v/v) with 10 mM ammonium formate and 0.1% formic acid, while mobile phase B was IPA/MeOH (80:20, v/v with 10 mM ammonium formate and 0.1% formic acid. The gradient was set as follows: 0-4 min 40% B; 4.1-6 min 40→60% B; 6.1-16 min 60→100% B; 16.1-22 min 100% B; 22.1-24 min 100→40% B; 24.1-30 min 40% B. Mass spectrometry analysis was made in positive ion mode with the following source parameters: spray voltage 3.5 kV, capillary temperature 320 °C, sheath gas 40 a.u., auxiliary gas 3 a.u., and S-lens RF 50. Data were acquired using a full-scan ddMS² Top 10 method with a total runtime of 30 min. Full-scan MS settings were: resolution 70,000, AGC target 1 × 10□, maximum IT 50 ms, and scan range m/z 200–1200. For ddMS², resolution 17,500, AGC target 1 × 10□, maximum IT 50 ms, and NCE 30 were applied. The injection volume was 3 µL.

Raw data acquired from LC-MS/MS analyses were processed using MS-DIAL software (version 5.25; RIKEN, Yokohama, Japan). The software was employed for peak detection, deconvolution of MS/MS spectra, compound identification, and alignment across all samples. Lipid species were identified by matching MS1 and MS2 data against internal libraries and public lipidomic databases integrated into the software. Quantitative values (nmol/mL) were derived by normalizing the integrated peak areas of each lipid species to the corresponding internal standard (same lipid class).

### AAV vectors design and organoid treatment

An AAV serotype testing panel (*in vitro* grade, PANEL-AAVS01, CMV-*EGFP*) was purchased from VectorBuilder. The rAAV6 *LIPA* vector was designed as follows: the cDNA of the human *LIPA* gene (Gene ID: 3988) fused to a C-terminal 3×HA tag^48^ was cloned into a standard rAAV2/6 vector backbone under the control of the human α1-antitrypsin (hAAT) promoter and an optimized *HBB2* intron^49^. All reagents and detailed sequence information are available upon request. Recombinant single-stranded rAAV2/6 particles were produced in HEK293 cells using an adenovirus-free triple transfection method and purified by single-step affinity chromatography (AVB Sepharose; GE Healthcare, Chicago, IL, USA). The final viral preparation was formulated in sterile phosphate-buffered saline containing 0.001% Pluronic F-68 (Sigma-Aldrich, St. Louis, MO, USA) and stored at −80 °C. At day 15, HLOs were collected using Cell Recovery Solution (Corning, 354253), and pelleted by gentle centrifugation (150g, 5 minutes). The pellet was resuspended in 300 µL of HCM and plated in suspension in a low-attachment 24-well plate. A concentration of 5.0 x 10^12^ (vg/mL) of AAV suspension were added per well and organoids were incubated overnight at 37 °C in 5% CO□. The following day, organoids were washed three times with PBS to remove residual virus and re-embedded in BME drops.

### Quantification and statistical analysis

Lipidomics and scRNA-seq data were analyzed using R, version 4.2.0. The mean Hnf4a fluorescence intensity was measured in individual DAPI positive nuclei. Nuclei were classified as Hnf4a positive using a fixed intensity threshold applied across all samples. The density of Hnf4a positive nuclei was calculated as the number of positive nuclei per mm² of analyzed tissue area. All statistical analyses were performed using GraphPad Prism 10. The association of a normally distributed variable between two groups of interest was assessed using Student’s t test. When more than 2 groups were considered, a one-way ANOVA was used to check for statistical association. Lastly, a two-way ANOVA was conducted to examine the influence of two independent variables on a dependent variable. Dunnett’s post-hoc test was used when one condition was taken as the control; otherwise, Tukey’s test was applied. Quantitative data are shown as mean ± standard deviation (SD) and are considered statistically significant when p < 0.05. In all the figures p-values are summarized as follow: ns = not significant, *p < 0.05, ** p < 0.01, *** p < 0.001, **** p < 0.0001. Unless otherwise indicated, all quantitative analyses were performed on samples generated from independent HLO differentiation experiments, with each differentiation considered a biological replicate.

## DATA AVAILABILITY STATEMENT

All data supporting the findings of this study are available from the corresponding author upon reasonable request.

## AKNOWLEDGEMENTS

G.S. is funded by grants from: Telethon (GGP20031) and Italian Ministry of University and Research and European Union under NextGenerationEU “Progetti di rilevante interesse nazionale” (PRIN): PRIN prot. no.2022PWKZXE; PNRR M4C2 INV 1.1, PRIN 2022 prot. no. P2022A9J9L; NRRP NextGenerationEU Project CN00000041-National Center for Gene Therapy and Drugs based on RNA Technology; AIRC (Start-Up 2020 – ID. 24322 project – PI S.G.). M.A. is funded by the French National Research Agency (grants: NEEDED ANR-24-CE18-3712-01). This study was partially funded by the Italian Ministry of University and Research (MUR) program, “Departments of Excellence 2023–2027”, the AGING Project—Department of Translational Medicine, University of Piemonte Orientale, and is part of the project, P2022E3BTH-CUP C53D23007600001, which received funding from NextGeneration EU-MUR M4C2–PRIN 2022 PNRR. G.D.S. is funded by the Fondazione AIRC IG grant 30570, the Fondazione AIRC Special Program Molecular Clinical Oncology “5 per mille” 22759, Worldwide Cancer Research (grant 24-0361), the NRRP NextGenerationEU Project CN00000041-National Center for Gene Therapy and Drugs based on RNA Technology, the Italian University and Research Ministry PRIN P2022ZWY8H and the PRIN 2022XBYNJP. R.B. was supported by an AIRC post-doc fellowship for Italy (ID: 29960).

The authors thank all the members of ADM lab at ICGEB for insightful discussions and Marco Bestagno from ICGEB for the technical support for the FACS analysis. Furthermore, we would like to thank Brassier Anaïs for helping in procuring patient samples. The figures in this work were created with BioRender (https://BioRender.com).

## AUTHOR CONTRIBUTIONS

Conceptualization, D.S. and G.S.; Investigation, D.S., C.D.R., B.A., S.V., R.B., K.H., M.L., A.A., A. Marfoglia, A. Mattivi, M. Manfredi.; writing—original draft, D.S., M.A. and G.S.; writing—review & editing, D.S. and G.S.; funding acquisition, G.S.; resources, L.C., M. Amendola., M.L., N.R., M. Mastronardi., S.P., N.d.M, D.B., P.D. and F.Z.; supervision, L.C., L.F., M. Amendola, C.T., D.C., G.D.S. and G.S.

## ETHICS APPROVAL AND CONSENT TO PARTICIPATE

The generation of organotypic cultures was approved by the ethical committee of the region Friuli Venezia Giulia (“comitato etico unico regionale”) protocol n. 0033969, 26/08/2024. Informed written consent was obtained from all subjects involved in the study.

---

**Supplementary Fig. 1:**
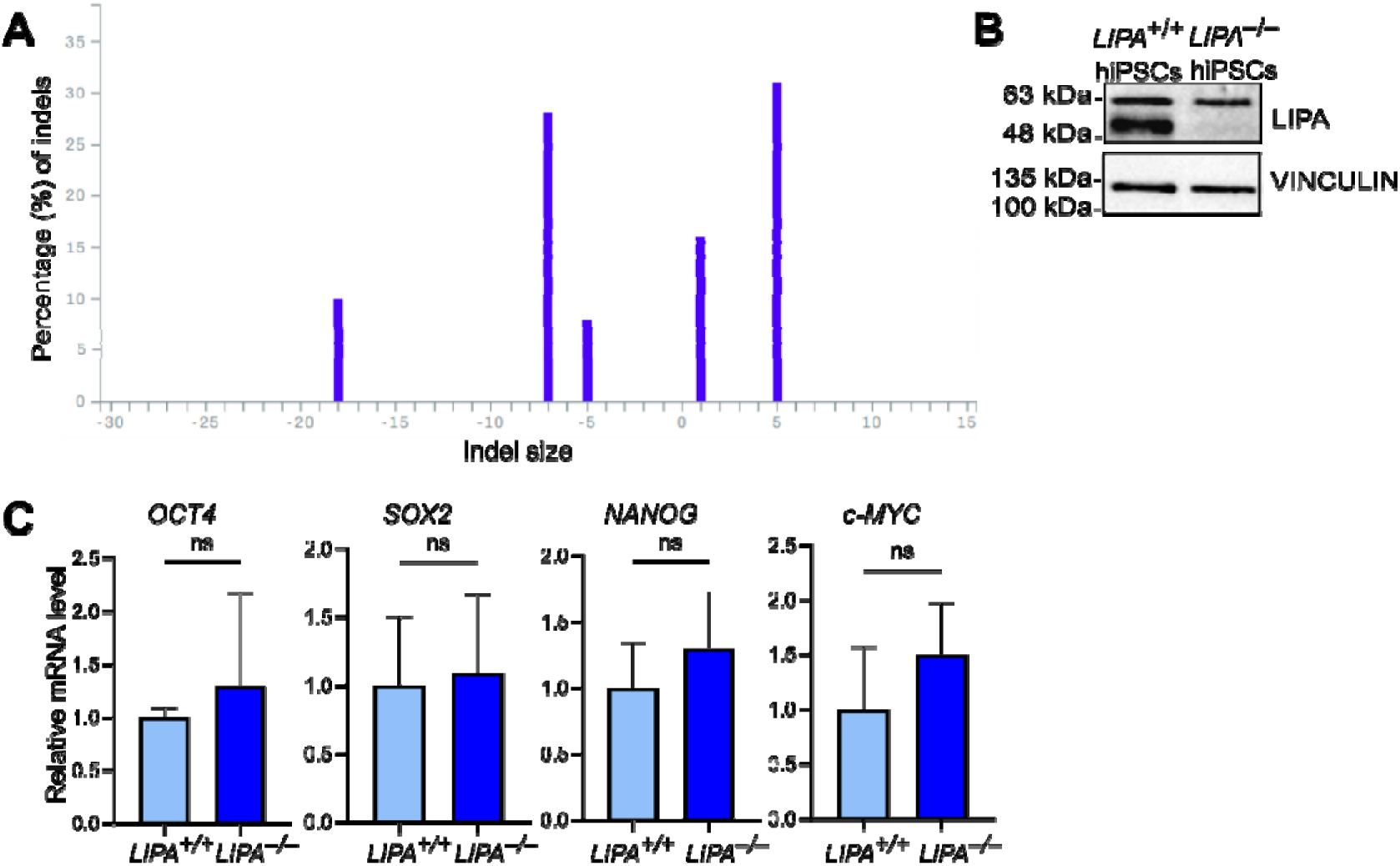
**A** Sanger trace deconvolution of the edited LIPA locus showing the distribution of indel sizes generated by CRISPR–Cas9. **B** Western blot analysis showing the reduction of LIPA protein levels following CRISPR/Cas9 gene editing in *LIPA^+/+^*and *LIPA^-/-^*iPSCs. **C** Quantitative RT–PCR analysis of pluripotency markers in *LIPA^+/+^* and *LIPA^-/-^* iPSCs. Results represent mean ± SD; n = 3. Student’s t-test was applied: ns = not significant.

**Supplementary Fig. 2:**
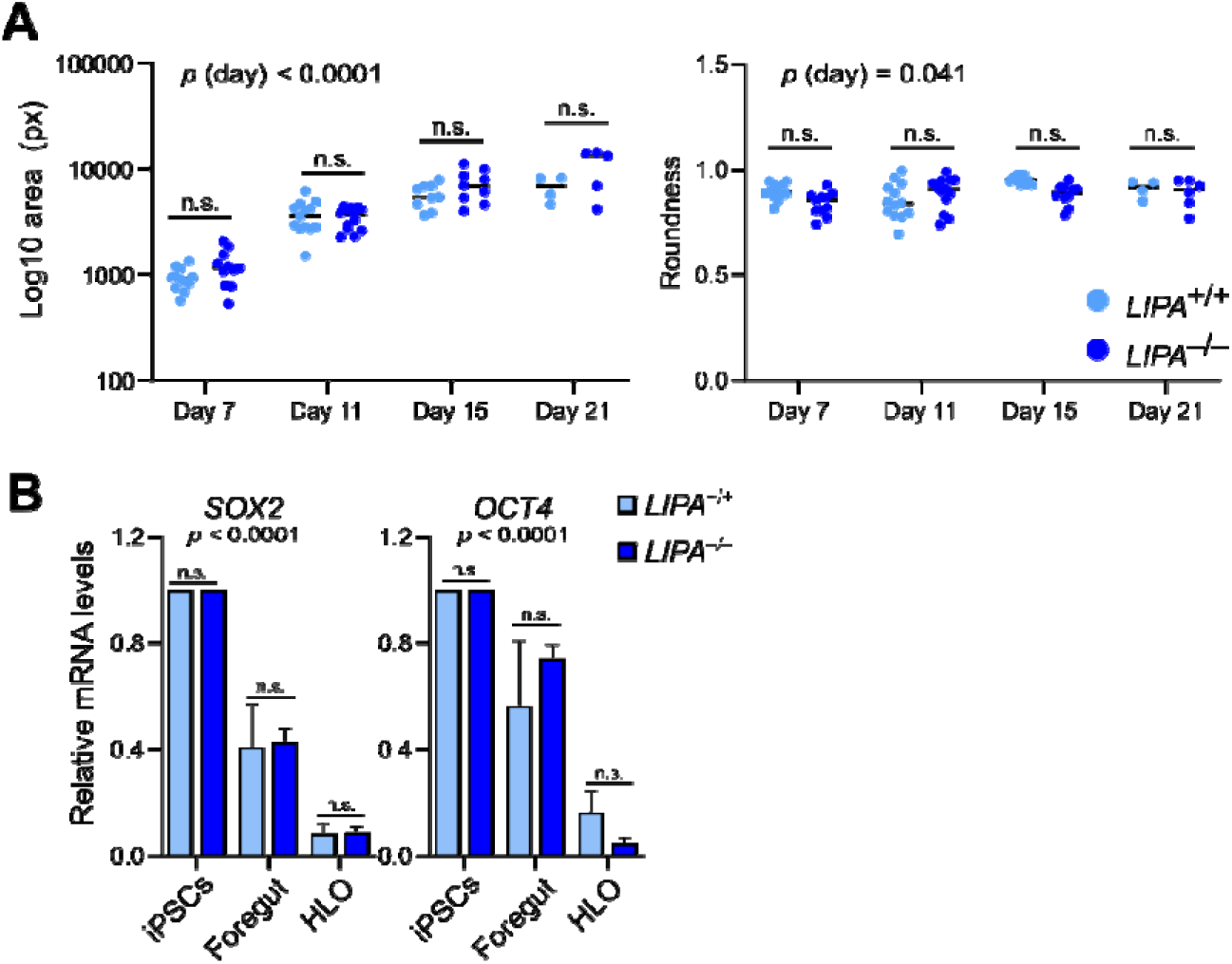
**A** Quantification of shape/morphological descriptors in *LIPA^+/+^* and *LIPA^-/-^*HLOs at day 7, 11, 15 and 21. **B** Quantitative RT–PCR analysis of pluripotency markers in *LIPA^+/+^* and *LIPA^-/-^* HLOs. Results represent mean ± SD; n = 3. Student’s t-test was applied: ns = not significant.

**Supplementary Fig. 3:**
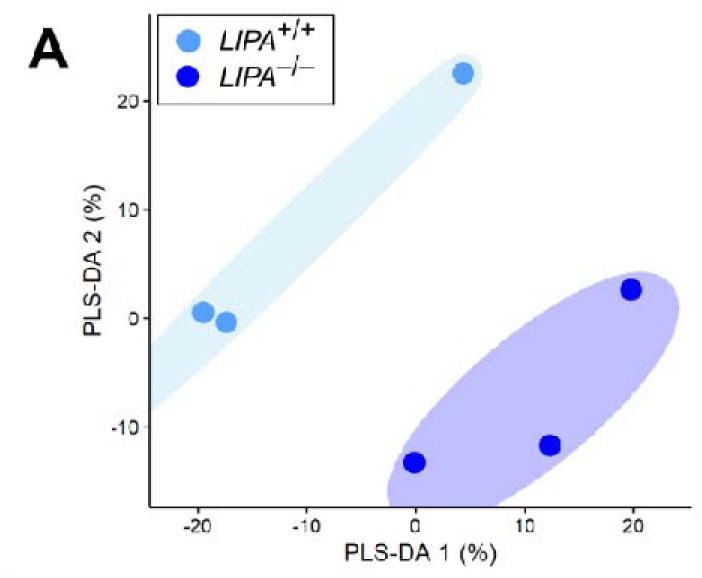
**A** Partial least squares discriminant analysis (PLS-DA) score plot showing the separation of *LIPA^+/+^* and *LIPA^-/-^* HLO samples based on their lipidomic profiles.

**Supplementary Fig. 4:**
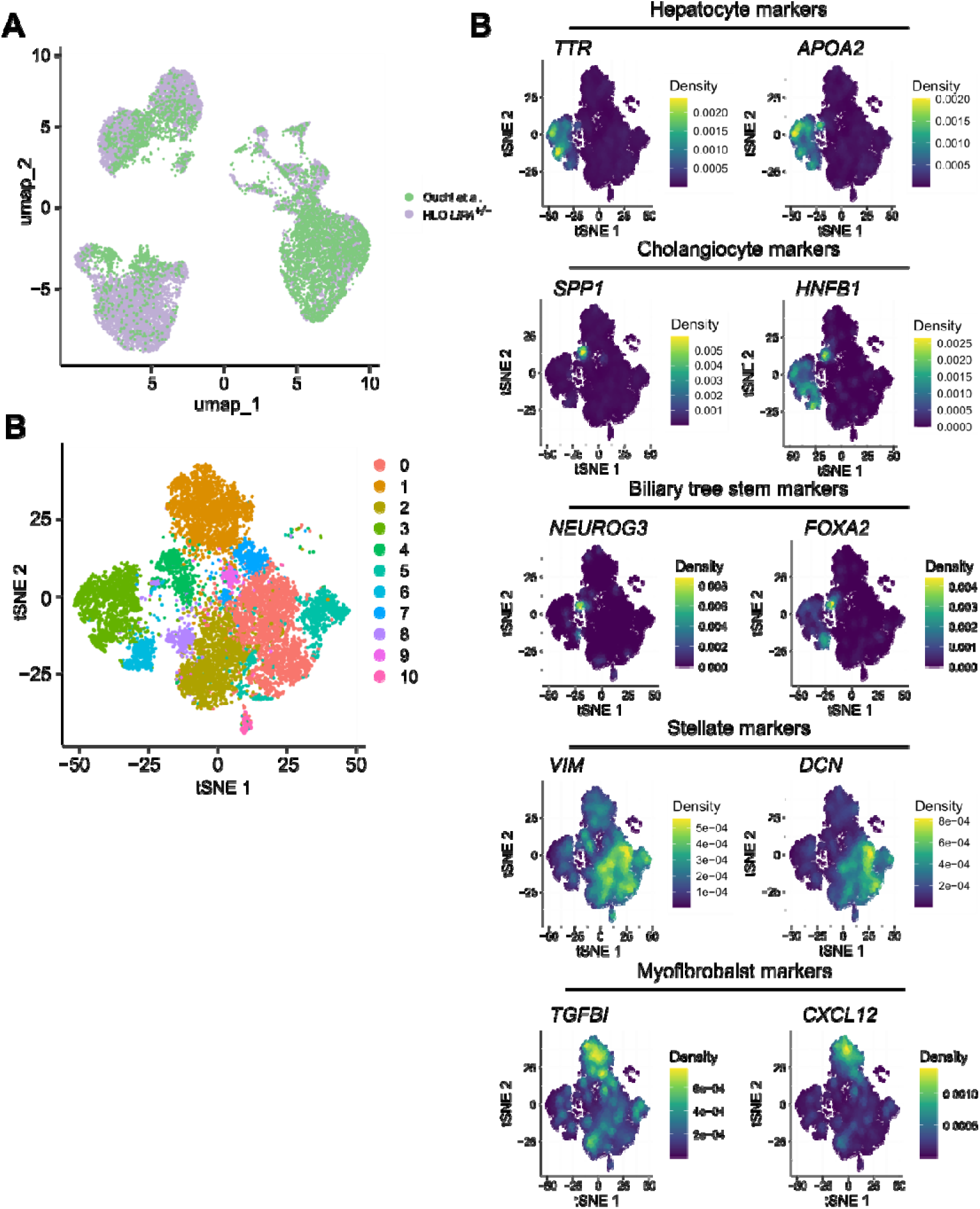
**A** UMAP visualization of HLOs from the present study and from Ouchi *et al*., 2019 shown by dataset source. **B** UMAP plot showing cell clusters identified in HLOs after quality control. **C** Density plots displaying the expression of representative cluster marker genes in HLOs.

**Supplementary Fig. 5:**
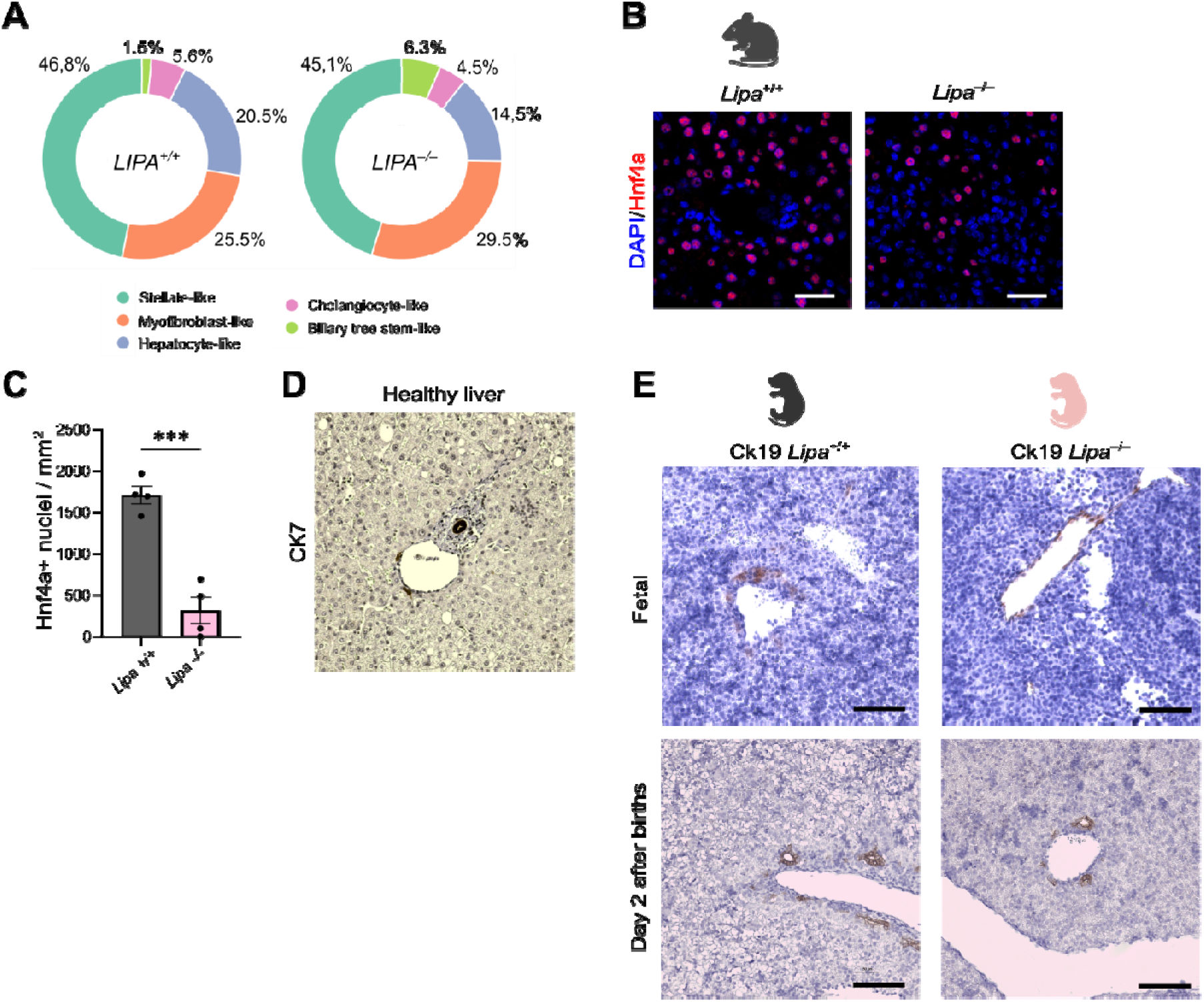
**A** Pie chart showing the relative proportion of cells in *LIPA^+/+^* and *LIPA^-/-^*HLOs. **B** Representative IF staining of Hnf4a in livers from *Lipa^+/+^* and *Lipa^-/-^* mice. **C** Quantification of Hnf4a positive nuclei in *Lipa^+/+^* and *Lipa^-/-^* liver sections. Data are presented as the density of Hnf4a positive nuclei (cells/mm²). Student’s t-test was applied: *** *p*□<□0.001. **D** IHC staining of CK7 in liver from non-WD donor. **E** Representative IHC staining of CK19 in livers from fetal and day 2 *Lipa^+/+^* and *Lipa^-/-^* mice. The scale bar represents 100 μm.

**Supplementary Fig. 6:**
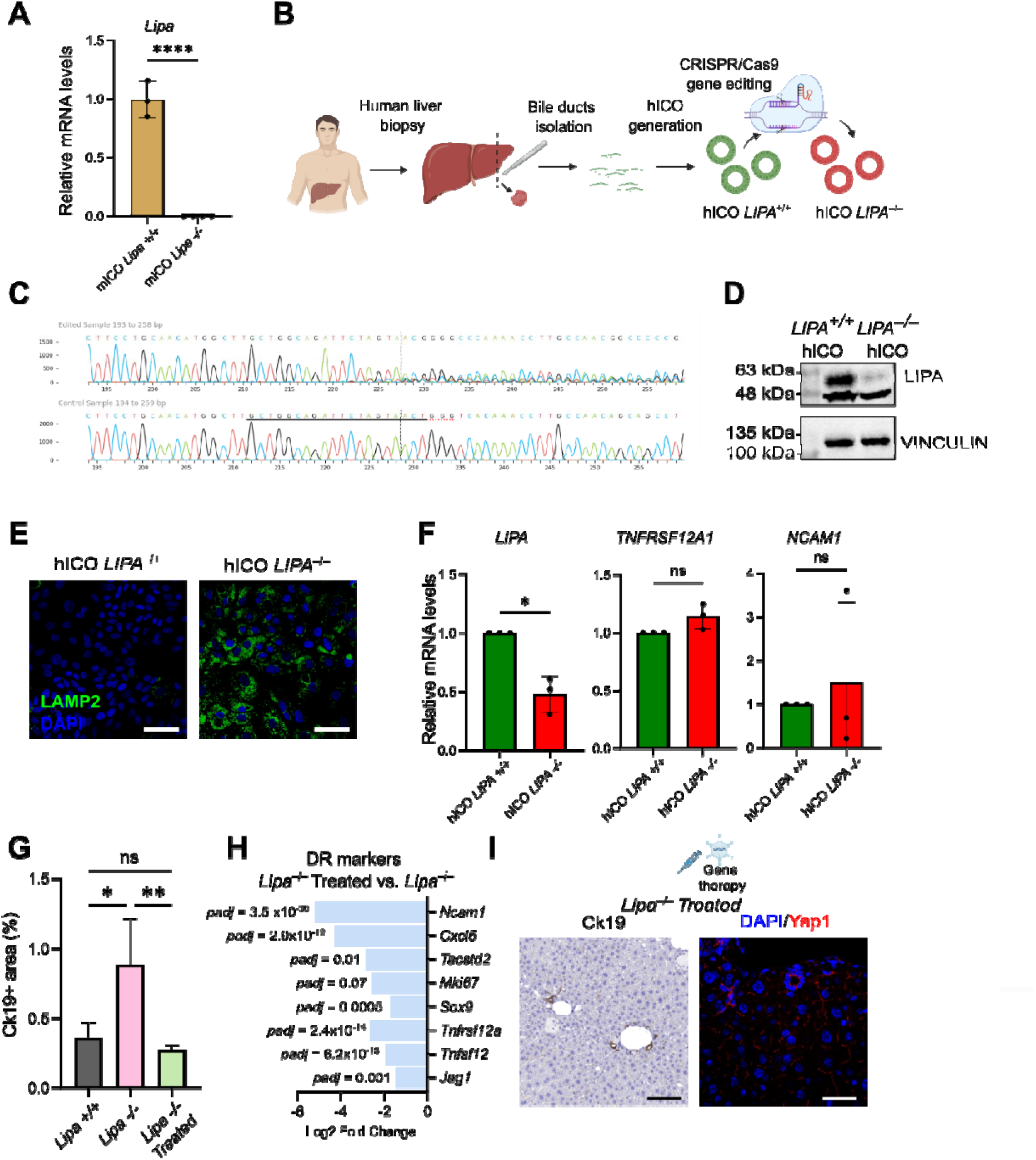
**A** Quantitative RT–PCR analysis of *Lipa* gene in *Lipa^+/+^* and *Lipa^-/-^* mICO. Results represent mean ± SD; n = 3. Student’s t-test was applied: **** *p*□<□0.0001. **B** Schematic representation of the *LIPA^-/-^* human cholangiocyte organoids (hICO) generation via CRISPR/Cas9 gene editing. **C** Representative Sanger sequencing chromatograms confirming the successful editing of the *LIPA* locus in hICO. **D** Western blot showing a reduction of the LIPA protein after gene editing in hICO. **E** LAMP2 immunofluorescent staining in *LIPA^+/+^*and *LIPA^-/-^* hICOs. The scalebar represents 50 μm. **F** Quantitative RT–PCR analysis of *LIPA* and DR markers in *LIPA^+/+^* and *LIPA^-/-^*hICO. Results represent mean ± SD; n = 3. Student’s t-test was applied: * *p* ≤ 0.05, ns = not significant. **G** Ck19 area quantification. One-way ANOVA was applied followed by Tukey post-hoc test. ns = not significant, * *p*□≤□0.05, ** *p*□≤□0.01. **H** Downregulated DR markers in whole liver of *Lipa^-/-^* Treated vs. *Lipa^-/-^* mice from Laurent et al. **I** Representative IHC staining of CK19 (lest panel) and IF staining of Yap1 and Hnf4a in livers from *Lipa^-/-^* treated mice (middle and right panel respectively). Representative staining images supporting the data presented in Figures 5C and 5F. The scale bar represents 100 μm.

**Supplementary Fig. 7:**
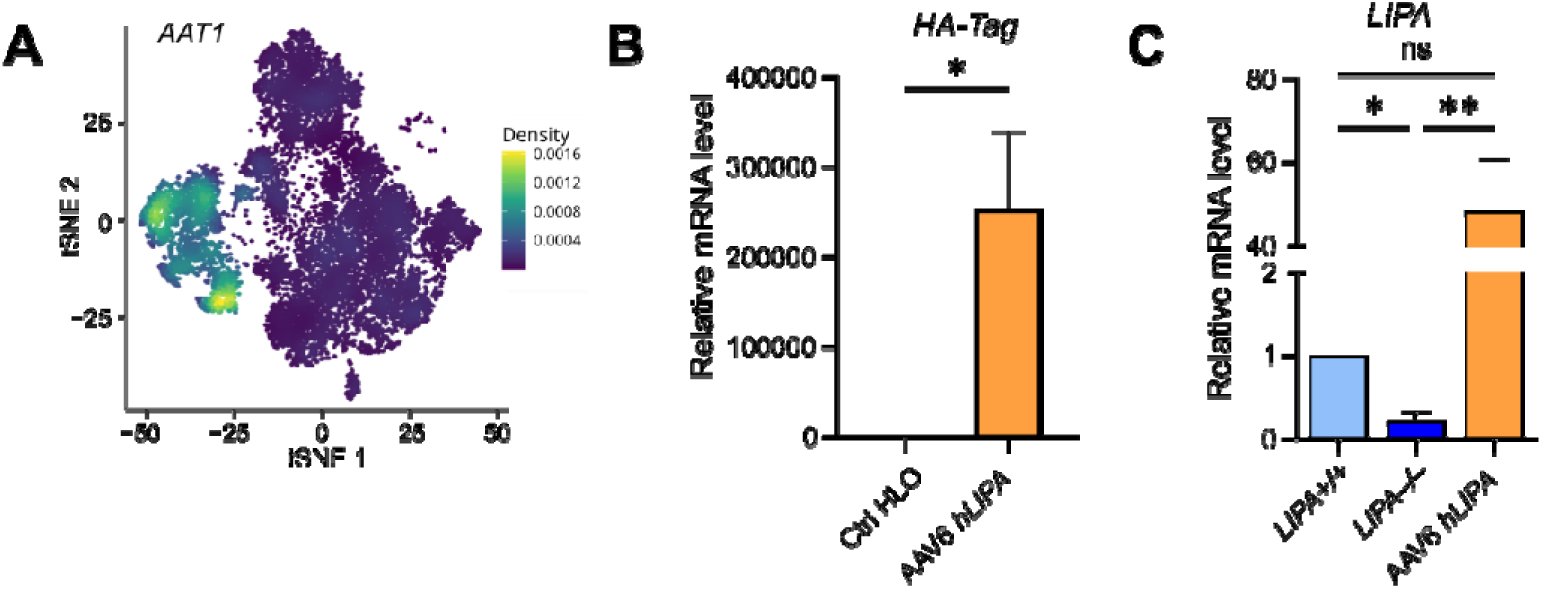
**A** Density plots displaying the expression of *AAT1* in HLOs. **B** Quantitative RT–PCR analysis of *HA-tag* in AAV6-*hLIPA* transduced HLO and in controls. Results represent mean ± SD; n = 4. Student’s t-test was applied: * *p*□≤□0.05. **C** Quantitative RT–PCR analysis of *LIPA* in *LIPA^+/+^*, *LIPA^-/-^* and AAV6-*hLIPA* HLO. Results represent mean ± SD; n = 4-6 independent differentiation experiments. One way ANOVA was applied with Tukey post hoc test: * *p*□≤□0.05, ** *p*□≤□0.01.

